# Diversity of ribosomes at the level of rRNA variation associated with human health and disease

**DOI:** 10.1101/2023.01.30.526360

**Authors:** Daphna Rothschild, Teodorus Theo Susanto, Xin Sui, Jeffrey P. Spence, Ramya Rangan, Naomi R. Genuth, Nasa Sinnott-Armstrong, Xiao Wang, Jonathan K. Pritchard, Maria Barna

## Abstract

Ribosomal DNA and RNA (rDNA and rRNA) sequences are usually discarded from sequencing analyses. But with hundreds of copies of rDNA genes it is unknown whether they possess sequence variations that form different types of ribosomes that affect human physiology and disease. Here, we developed an algorithm for variant-calling between paralog genes (termed RGA) and compared rDNA variations found in short- and long-read sequencing data from the 1,000 Genomes Project (1KGP) and Genome In A Bottle (GIAB). We additionally developed a novel protocol for long-read sequencing full-length rRNA (RIBO-RT) from actively translating ribosomes. Our analyses identified hundreds of rDNA variants, most of which, surprisingly, are short insertion-deletions (indels) and dozens of highly abundant rRNA variants that are incorporated into translationally active ribosomes. To visualize variant ribosomes at the single cell level, we developed an *in-situ* rRNA sequencing method (SWITCH-seq) which revealed that variants are co-expressed within individual cells. Strikingly, by analyzing rDNA, we found that variants assemble into distinct ribosome subtypes. We discovered that these subtypes acquire different rRNA structures by successfully employing dimethyl sulfate (DMS) probing of full length rRNA. With this atlas we investigated rRNA variation changes across human tissues and cancer types. This revealed tissue-specific rRNA subtype expression in endoderm/ectoderm-derived tissues. In cancer, low abundant rRNA variants can become highly expressed, which suggests the presence of cancer-specific ribosomes. Together, this study identifies and comprehensively characterizes the diversity of ribosomes at the level of rRNA variants which is dominated by indel variants, their chromosomal location and unique structure as well as the association of ribosome variation with tissue-specific biology and cancer.

## Introduction

The ribosome is a complex, ancient machine responsible for all protein synthesis, with a core ribosomal RNA (rRNA) structure that is conserved across all kingdoms of life. The primary transcript of the rRNA genes is the large 45S pre-rRNA which contains the 18S, 5.8S, and 28S rRNAs as well as transcribed spacer regions. In humans, rRNA genes are present in hundreds of ribosomal DNA (rDNA) copies in tandem repeats that are spread across multiple chromosomal loci ^1^. These high rDNA copy numbers are thought to be necessary to produce millions of ribosomes in each cell. Nevertheless, these hundreds of rDNA copies allow for sequence variation between copies, as was first noted in mice and humans almost 50 years ago ^2^.

It remains an outstanding challenge to understand whether ribosomes are different at the level of rRNA and how many ribosome subtypes may exist. For the last several decades we have therefore had a limited knowledge of fundamental differences in the translational machinery and limited insight beyond the textbook view of ribosome composition. A very significant study showed that rRNA sequence variations between different rDNA loci lead to novel rRNA functions, in *E.coli*, where sequence variations between the bacterial rDNA operons support growth under stress conditions ^3^. In particular, they found that naturally occurring rRNA sequence variation can modulate bacterial ribosome function, central aspects of gene expression regulation, and cellular physiology. This inspiring work raises the possibility that this is general and could be the case in humans. In humans, by examining short-read sequencing data of the 1,000 Genomes Project (1KGP) ^4^, two previous studies discovered hundreds of positions in rDNA bearing sequence variants ^5,6^. A major challenge faced by these former studies arises from limitations in short-read sequencing where there is the depletion of GC-rich sequences. Specifically certain regions within rDNA possess >80% GC content on average ^6–8^. This results in two types of problems in variant calling: (1) false negatives, caused by the inability to identify variants in regions with poor sequence coverage, and (2) false positives, caused by PCR errors in low-coverage regions. Moreover, these GC rich regions are highly repetitive which make short-read variant discovery tools inaccurate ^9^. As a result, the latter two previous studies ^5,6^ reported contradictory results likely because of these major caveats. Nevertheless, these studies did raise interest in investigating rRNA sequence variation, inspiring future studies. Interestingly, the copy number of the rDNA loci was also shown to be important for gene expression and cellular homeostasis ^10–13^. Moreover, rDNA copy number was associated with age and in cancer ^14–16^. In cancer, both rDNA copy number gain and loss have been reported in multiple studies ^17–21^. This calls for further investigation into whether rDNA copy number changes are coupled with sequence variations and if such changes affect human health.

Recently, long-read sequencing enabled the successful complete assembly of a human genome by the Telomere-2-Telomere (T2T) consortium, including positioning of rDNA copies in the five acrocentric chromosomes ^22^. Moreover in the mouse and Arabidopsis plant genomes, rDNA variants were grouped into haplotypes and a few rRNA variants were found expressed in tissues using short-reads ^5,23,24^. Yet in order to find low-frequency variations between full length rDNA paralog copies, new computational method development is necessary. Long-reads offer the possibility for distinguishing between paralog genes. However, existing common methods for long-read variant calling, such as DeepVariant ^25^ and Clair ^26^, are primarily designed for detecting variants in single-copy regions. For paralog genes where low-frequency variants exist between copies accurate variant calling is lacking ^9^. Moreover, it remains an open question if rRNA haplotypes are expressed from the human genome, what their abundances are, and if such variability is linked to human physiology. There too, long-read sequencing and analysis of full-length rRNA is necessary This highlights the need for new approaches to comprehensively characterize human ribosome diversity.

To address this need in the field, we devised an efficient novel computational algorithm to detect all variations between paralogs (termed RGA). Applied to the long-read 1,000 Genomes Project dataset, we discovered hundreds of rDNA sequence variations enriched with previously undiscovered insertion-deletions. We further developed a novel methodology to perform long-read sequencing on rRNA in actively translating ribosomes to identify variants (RIBO-RT). Using this method, we discovered that ribosomes have different subtypes with rRNA variants that are genomically encoded by rDNA clustered on distinct chromosomes. Additionally, using an *in-situ* rRNA sequencing platform that we developed (SWITCH-seq), we discovered that variants belonging to different rRNA subtypes are co-expressed in single cells. We then used structure probing coupled with long-read sequencing to find that 28S subtypes have different rRNA structures. Lastly, we found that these subtypes are differentially expressed in human tissues, and low abundance variants are elevated in certain cancers. Together, these results suggest that ribosomes with unique sequence variation may be used to modulate different cellular programs underlying human physiology and disease.

## Results

### Indels are the main variants of the human rDNA loci

How rDNA variation shapes the presence of unique ribosomes in the cell remains an important open question. Previous studies that analyzed the 1KGP dataset for discovery of rDNA variants reported discordant results. Parks et al. ^5^ reported hundreds of variants in both the 18S and the 28S, yet 75% of variants were not made publicly available, making a comparison to this dataset problematic. Nonetheless, Fan et al. ^6^ reported notable differences from Parks et al. by suggesting that the 18S has low variation, and also reporting many fewer variant positions in the 28S. Moreover, Parks et al. reported only 2.7% of variants being indels while Fan et al. reported 19.2% indels. Here, considering the limitations of short-reads in rDNA variant discovery and their inability to distinguish between rDNA paralogues we decided to re-evaluate the variants in the human rDNA genes.

Until recently, the 1KGP dataset included only short-read genome sequencing ^27^. Yet as of 2022, the 1KGP includes PacBio’s HiFi long-read sequencing for 30 individuals from diverse ancestral origins, which could serve as a better method for accurately calling variants. Here, we compared the rDNA variants captured by short- and long-reads from the same individuals and addressed the discrepancies between previous studies (**Figure 1A,B, Table S1**).

**Figure 1.**
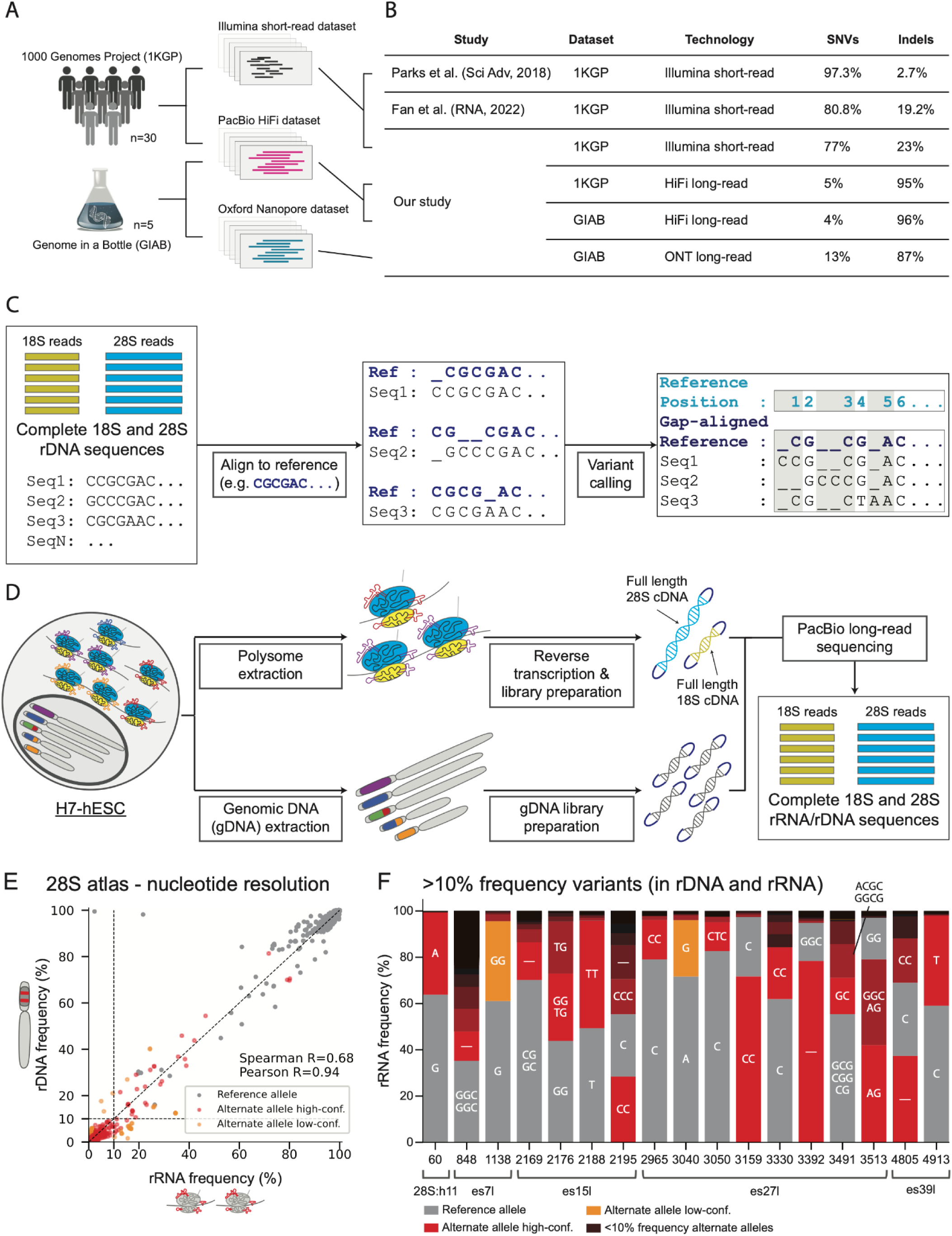
1,000 Genome Project (1KGP) and H7-human embryonic stem cells (hESCs) rRNA and rDNA variant extraction pipeline with high correlation between 28S rDNA and rRNA high abundant variant frequencies. A. Graphical illustration of the dataset analyzed consisting of 30 individuals from the 1000 genomes project (1KGP) with both short and long-read sequencing. B. Comparison of Single Nucleotide Variants (SNVs) and Insertion-deletions (Indels) across studies. C. Graphical illustration of the Reference Gap Alignment (RGA) alignment method used for variant discovery in 18S and 28S sequences D. 18S and 28S rDNA / rRNA sequence extraction pipeline from H7-hESC E. Scatter plot of 28S rRNA frequency (x-axis) and rDNA frequency (y-axis) for reference and alternate alleles. Alternate alleles are marked in red if their frequencies in rDNA and rRNA agree or in orange if they differ significantly. Reference alleles are marked in gray. A dashed black line indicates rRNA frequency equal to 10%. Spearman and Pearson correlations for rRNA frequency and rDNA frequency between alternate alleles alone are presented (calculated on variants, red dots alone). F. Stacked bar plots of allele frequencies at positions with variants with frequency >10% in both rRNA and rDNA. The nucleotide sequence matching the alleles are indicated inside the bar plots for variants with >10% allele frequency (‘-’ indicates deletion). The reference allele is indicated in gray and alternate alleles are indicated in color.

When analyzing the short-read data, we followed the pre-processing steps as performed in previous studies of marking duplicate reads which are suspected as PCR artifacts. This step discarded 97% of reads. However, it is unknown if duplicate reads are PCR biases given the high rDNA paralog copy numbers which highlights the limitation of short-read sequencing for rDNA variant discovery. Next, in order to call variants including rare variants which are not expected to follow germline variant frequencies in high paralogues rDNA copy numbers, we tested two common somatic variant calling methods for short-reads: LoFreq* ^28^ which was used by Parks et al. and Mutect2 ^29^. Specifically, Mutect2 was chosen instead of the germline variant caller used by Fan et al., because germline variant callers will not detect rare variants found between paralogues. Using the LoFreq* method which is known to be sensitive ^28^, we found 1,582 positions with variants compared to 861 positions with variants with Mutect2 (**Table S2-3**). Notably, both methods detected 23% indels, which is on par with the indel percentage reported by Fan et al.. Given the difference in the proportion of indel frequencies between the previous two studies, we compared variant quality scores of SNVs and indels (**Table S3** - Mutect2 FDR-corrected log 10 likelihood ratio score of variant existence). Here we found that indel variants were enriched with high confidence P-values (**Figure S1**, P-value < 10^-15^ comparing SNVs and indels likelihood ratio scores using Kolmogorov-Smirnov test for goodness of fit). Notably, tandem repeats and GC-rich sequences in the human genome were shown to be prone to chromosomal breakage and were found enriched in indels ^30^. Therefore, the 23% indel frequency derived solely from short-read data may underrepresent the true indel frequency in these samples. To test this, we next examined the HiFi long-read data from the same samples.

LoFreq* and Mutect2 did not work on the long-read sequencing data. In order to identify all positions with sequence variants, we developed a new computational method for acurate variant calling between paralogs which we term Reference Gap Alignment (RGA, see **Methods**). We align all reads against a common reference, and report all variants at a given position with respect to this reference (**Figure 1C, S2, Methods**). Our method reports at every position with respect to the 18S/28S reference if that position has a variant and calls its identity. The only parameters in our method are the standard pairwise alignment parameters: a mismatch penalty, a gap opening penalty, and a gap extension penalty (see **Methods** for more details). When benchmarking the global sequence alignment parameters, these resulted in similar indel proportions (**Table S4**). In agreement with previous studies ^5,6^, we found that rDNA is highly variable, yet using our method, we discovered that the vast majority of variants are short indels and not SNVs in all 30 samples. Specifically, when examining each reference position individually we found that on average 95% of variants are GC-rich indels (**Table S5**).

Since this indel proportion found with long-reads is markedly different from the results obtained with short-reads (**Figure 1B)**, we cross-validated our reported variants in three ways. As first indel validation, we decided to compare our results from our primary long-read sequencing technology, namely HiFi, with an alternative, Oxford Nanopore (ONT). Importantly, ONT is much more error prone compared to HiFi, with an estimated 13% error rate in ONT compared to 0.1% error rate in HiFi ^31–33^. Since 1KGP long-read sequencing was only performed on HiFi, we tested this using the Genome In A Bottle (GIAB) dataset, which consists of two trio families, where both HiFi and ONT were performed on the same samples. Here, when examining the HiFi dataset of GIAB, in agreement with the 1KGP HiFi results, we discovered that 96% of variants are indels (**Table S6**, **Methods**). When using the ONT dataset as a validation dataset, 81% of variants found in HiFi were replicated in the ONT dataset (**Table S7, Methods**). Notably, the variants that were not identified by ONT consisted of insertions and SNVs but not deletions (**Table S8**). Additionally, after retaining variants found at frequencies above the ONT error rate, 87% of found variants were indels (**Table S7** filtering variants with allele frequency smaller than 0.13). As another method to cross-validate our RGA variant caller, we examined variant frequencies in the family members of the GIAB trio dataset. We compared variant frequencies for variants that appeared only in the child, in a single parent, or in both parents. Here we expected that high abundant variants would be inherited and indeed all variants with frequency greater than 2% were found in at least one of the parents. This held true for both for SNVs and indels (**Figure S3**).

Finally, we tested if short-reads contain reads with indel variants that are not identified by short-read variant callers. To do this, we used the indels identified with long-reads by the RGA method as a reference of existing sequence variations and mapped the short-reads from the 1KGP to this reference (**Methods**). This directly tests if the variants found in the long-reads are also detected in the short-reads. Notably, mapping variants is different from de-novo variant discovery since here we count short-reads which perfectly match the indels as opposed to variant calling tools which are reference free and need to appear in high enough frequency to be discovered. Surprisingly, despite estimated low coverage of the rDNA loci we could detect 928 nucleotide variants which included 896 indels and 32 SNVs (**Methods, Table S9**). Previous variant callers which have analyzed the 1KGP were not able to identify most indels, likely because they are too low in abundance (LoFreq*, Mutect2 and DNAseq pipeline Sentieon, Release 201911). Our results show that short-reads can be mapped to indels given a reference of variants identified with long-read sequencing.

Our results highlight that previous studies which used short-read sequencing missed the majority of the variants found in the rDNA loci. Most of our identified rDNA variants were found at GC-rich regions which are depleted from short-read sequencing. We conclude that indels are the main variants in rDNA both between and within individuals. These findings also highlight the need to curate a reference of rDNA variants and the importance of our accurate variant calling method together with long-read sequencing in confidently assigning rDNA variants.

### An atlas of 18S and 28S human rDNA variants validated in rRNA of translating ribosomes and single cell microscopy

It is unknown if the rDNA copies with sequence variants found in the human genome are transcribed and are found in functionally translating ribosomes. With no human reference of different rRNA subtypes, studies performing RNA sequencing completely ignore rRNA variants which limits our understanding of the contribution of rRNA to human physiology and disease.

Sequencing of rRNA has been historically technically challenging ^6,34^. Here, in addition to our computational RGA variant discovery method, by optimizing long-read sequencing, we have successfully developed an experimental method for full length rRNA sequencing of 18S and 28S from translating ribosomes which we named RIBO-RT. In order to extract rRNA from translationally active ribosomes, we first employed sucrose gradient fractionation wherein ribosomes can be separated into free ribosomal subunits and translationally active ribosomes, which contain one or more ribosomes bound to the same mRNA. We extracted RNA from translating ribosomes-containing fractions (**Figure S4**), performed reverse transcription in denaturing conditions, and then sequenced complete 18S and 28S rRNA by HiFi long-read sequencing (**Figure 1D**, **Methods**). We selected a human embryonic stem cell line (H7-hESC) and using long-read sequencing, we sequenced the 18S and 28S from both its rDNA and rRNA (**Figure 1D**). Here, we obtained 58,495 sequences of the 18S and 14,430 sequences of the 28S rRNAs from translating ribosomes (**Methods**). With this approach and our variant discovery method, we were able to coherently characterize the 18S and 28S H7-hESC rRNA variants. Most importantly, since rRNA is known to be heavily modified ^35^, our strategy of matching rRNA to genomic rDNA from the same cell enables us to distinguish modifications or sequencing errors from true sequence variants belonging to different rDNA alleles.

In agreement with our 1KGP and GIAB rDNA results, we found that the H7-hESC rDNA is highly variable and is enriched with indels. Moreover, 96% and 84% of the hESC variants are also found in the 1KGP genomes projects and GIAB datasets respectively. This is generally concordant with expected rates of replication based on these small sample sizes of the 1KGP and GIAB datasets. Moreover, we find high agreement in the frequency of variants between the hESC to other datasets (**Figure S5**). Additionally, rDNA variants are transcribed into functionally translating ribosomes as they are present in polysome fractions (**Figure S6**). Specifically, we found 270 positions with variants in the 18S and 858 positions with variants in the 28S, corresponding to about one variant every six rRNA positions (**Figure S6-7**). When comparing monosome to polysome fractions we observed high correlation in variant frequencies between fractions (**Figure S8, Table S10**). Additionally, we long read sequenced the 18S and 28S from translating ribosomes from an additional commonly used cell-line, K562 using the same extraction protocol as described in the methods (**Figure S9**). We found that 95% of the H7-hESC variants are also found in the tested human cell line (**Figure S10, Table S15**). Most variants (59%) are found in expansion segment (ES) regions (**Figure S11**, ES/non-ES regions are annotated). These regions vary in sequence both within and among different species, nearly doubling the eukaryotic rRNA sequence relative to that of prokaryotes ^36^. ESs have recently been shown to bind ribosome associated proteins and transcripts, yet their functions remain poorly understood ^36–40^.

Most importantly, with accurate variant calling and full coverage of the underlying rDNA and transcribed rRNA, we can measure the frequency of each variant between the rDNA copies and rRNA expression levels. We distinguish possible modifications or sequencing errors from certain sequence variants belonging to different rDNA alleles by calling variants with similar frequencies measured in rDNA and rRNA as high-confidence alternate allele variants while those significantly deviating between their rDNA and rRNA frequencies as low-confidence alternate alleles (**Figure 1E, Table S11,** Fisher’s Exact Test measuring association between rDNA and rRNA alternate allele frequencies - high-confidence alternate alleles are marked in red, low-confidence alternate alleles are marked in orange). Surprisingly, we did not identify any abundant high-confidence variants within the 18S. This suggests that rRNA variation is not tolerated within the small ribosome subunit. While most variants have low abundance, in the 28S we found 23 variants at 17 positions with frequencies above 10% in both rRNA and rDNA (**Figure 1E,F, Table S12, Figure S7** annotated positions, **S8C** minor allele frequency with dashed-like marking at 10% frequency). Notably, for the 28S high abundance variants, there is very good agreement between variant frequencies in rRNA and rDNA (**Figure 1E**, R=0.93 Pearson correlation, correlating all variants colored in red and orange).

Next, we focused on the 28S high abundance variants. For most positions, the RNA45S5 reference allele is the major allele found in the sequenced H7-hESC line (**Figure 1F**, reference allele in gray, **Extended Data 1-2 nucleotide atlas**). Yet some alternate alleles were more abundant than the 28S RNA45S5 reference alleles (**Figure 1F**, gray for the reference allele and red for variants). Notably, high abundant variants are only located in 4 ES regions (es7l, es15l, es27l and es39l) and one non-ES region, helix 28S:h11 (**Figure 1F**). Moreover, in the es7l, es15l and es27l regions we observed that variants can be grouped and characterized by indels of GGC in tandem-repeats. GGX tandem-repeats were recently suggested capable of forming G-quadruplex structures ^41^ while other works suggested that such repeats can form other higher order structures ^42,43^. While the function of these ESs is largely unknown, a growing body of research supports various roles in translation regulation. For example, es27l has been shown to be important for control of translation fidelity and binding of ribosome-associated proteins for several processes, such as initiator methionine cleavage from the nascent polypeptide ^44^ or acetylation of nascent polypeptides ^45^. Moreover, es39l interacts with the Signal Recognition Particle (SRP), which identifies the signal sequence on nascent polypeptides emerging from the translating ribosome ^46^. Interestingly, the most abundant variant in the non-ES helices, a G to A substitution at position 60 in 28S:h11, is considered unique to humans. The alternative allele, A, is the reference allele for other mammals including chimpanzees ^47,48^.

For these aforementioned highly abundant variants, a strong correlation between the frequency of a variant’s occurrence among rDNA copies and its expression levels in rRNA indicates both the authenticity of these sequence variants and their likely co-expression within individual cells. To explore this hypothesis, we developed a template-switching-based *in-situ* sequencing method (SWITCH-seq) to visualize variant ribosomes in individual HeLa cells (**Figure 2A**). This approach involved designing a reverse transcription primer to target constant non-variable regions downstream of the selected rRNA variant regions. Specifically, we selected regions for which we could design a primer for each of the ES regions with abundant variants (**Methods, Table S13**). The process of SWITCH-seq begins with performing reverse transcription on fixed HeLa cells, wherein a known sequence of choice is attached to the 3′ end of cDNA (template switching) (**Figure 2A**, **Methods**). This step incorporates the variant of interest into the cDNA, which is subsequently amplified into *in-situ* cDNA amplicons through enzymatic circularization and rolling circle amplification. These amplicons encapsulating the rRNA variants are then probed into a hydrogel network for cyclic imaging using a confocal microscope (**Methods**). We conducted multiple rounds of imaging which capture two bases before the variants, and the consecutive base, where we expect to find the variants (**Figure 2B, S12**). As predicted, we successfully observed both the reference and alternate alleles (**Figure 2B,C, S12**). Furthermore, the frequencies of the reference and variant alleles corresponded with their frequencies in the H7-hESC samples (**Table S14**). We conclude that rRNA variants observed at high frequency in the H7-hESC rRNA and rDNA and in the rDNA across the 1KGP and GIAB samples form ribosomes that are co-expressed in individual cells that can be visualized at single cell resolution.

**Figure 2.**
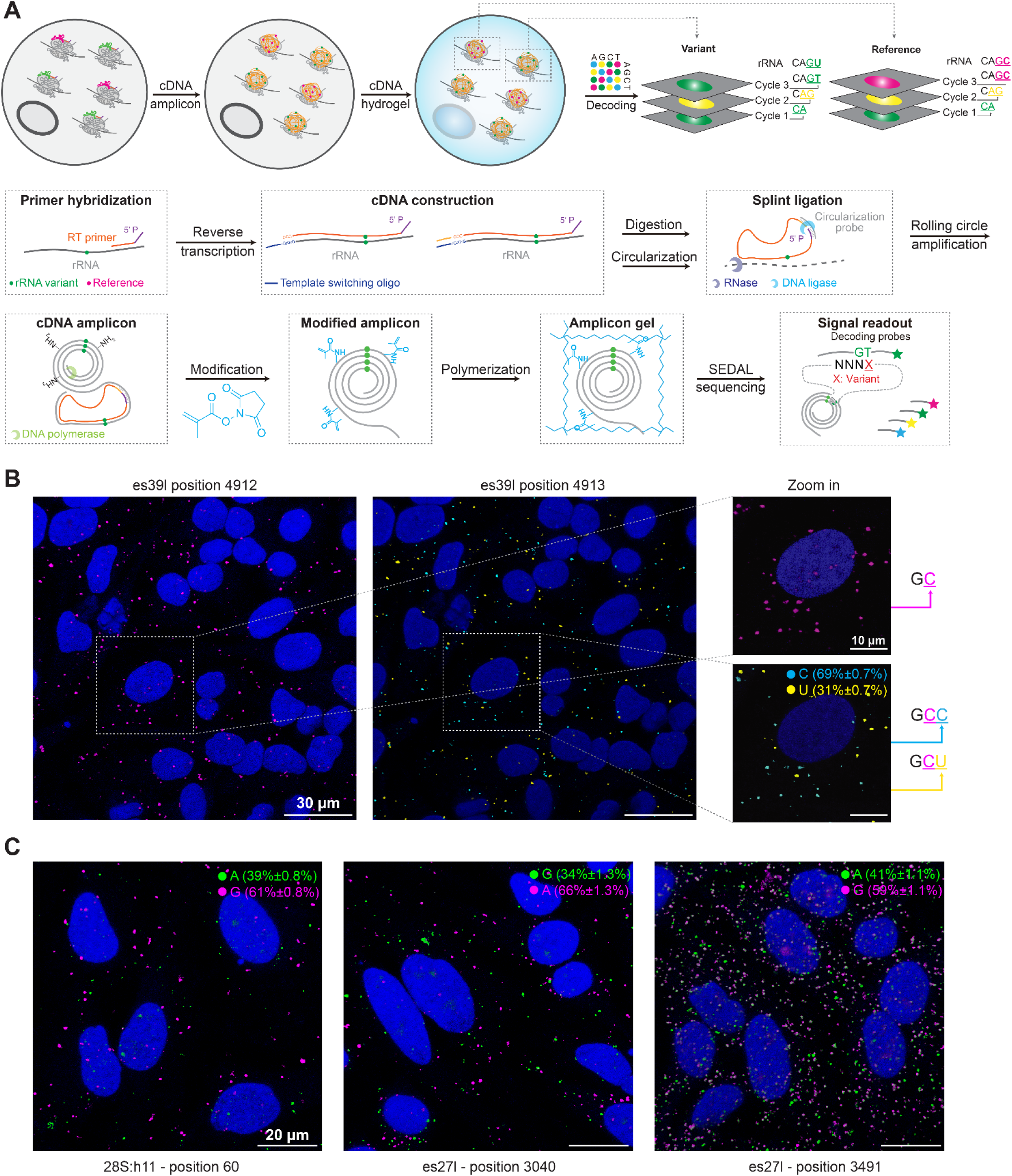
rRNA variants are co-expressed in individual cells as visualized by *in-situ* sequencing. A. Graphical illustration of SWITCH-seq pipeline. B. Two rounds of representative fluorescent *in situ* sequencing images of HeLa cells (DAPI staining in blue) are presented for the es39l-probed region. We identified a non-variable base C (magenta) at position 4912. At position 4913, two alternative sequences were revealed: the known reference sequence C (cyan) and the alternative variant U (yellow). Data shown as mean percentage ± standard deviation. n = 4 images. C. Representative fluorescent images of HeLa cells (DAPI staining in blue) showcase 3 highly abundant rRNA variants. The positions of the variants are indicated at the bottom of the images, while the reference (magenta) and alternative alleles (green) are indicated at the top, along with their respective rRNA frequencies. Data shown as mean percentage ± standard deviation. n = 4 images per allele.

As an important resource for studying human rRNA variations, we create the first comprehensive atlas of all H7-ESC rRNA 18S and 28S rRNA variants at different resolutions from nucleotide variants to gene 28S haplotype-groups that we later describe as separate subtypes (**Figure S11, Table S17-18** for region annotation and atlas building is described in the **Methods**).

### 28S variants assemble to distinct ribosome subtypes

Since we successfully obtained full length 18S and 28S rRNA with variations, we address the outstanding question of whether variations lead to the formation of different ribosome subtypes. Here we focused on the 28S since 18S variants appeared at low frequency. For the 28S, we found high agreement between rRNA and rDNA variant frequency so we first asked which rDNA variants are co-located on the same 28S rDNA copy. To do this, we calculated the correlation coefficient (Pearson’s r^2^) between positions across all 28S H7-hESC rDNA sequences. This is analogous to measuring the linkage disequilibrium (LD) coefficient in population genetics, though across paralogous copies within a single genome rather than across individuals in a population. Notably, we found low global LD structure between highly abundant rDNA variants (**Figure 3A** showing LD for rDNA positions with found rRNA frequency > 10%), supporting recent findings indicating high rates of nonallelic gene conversion across the acrocentric chromosomes ^49^. The highest LD (r^2^ > 0.2) was found between the es27l to all other regions. Comparing different regions, we found LD between four regions: 28S:h11, es15l, es27l and es39l, where in each region we identified a position with higher linkage to the other three regions (**Figure 3A**, four positions are annotated, position 60 being 28S:h11, with higher linkage to other positions). By considering the variants at this subset of positions, we found a total of 21 different haplotypes both in rDNA and rRNA (**Figure S13**). For testing if haplotypes can be considered as different 28S subtype variants, we further analyzed two independent long-read DNA datasets: (1) the fully assembled genome from the T2T with 219 rDNA copies with their chromosome location ^22^, and (2) the GIAB HiFi dataset ^50^. In agreement with the H7-hESC results, 3 out of the 4 positions with higher linkage to other positions in the H7-hESC had higher LD in the GIAB dataset (**Figure 3B, S14**). Since these three variants are linked to variants at other positions, we define the haplotypes formed by positions 60, 3513 and 4913, belonging to regions 28S:h11, es27l and es39l respectively, as different 28S haplotypes (**Figure 3C**, **Extended Data 7 28S atlas**).

**Figure 3.**
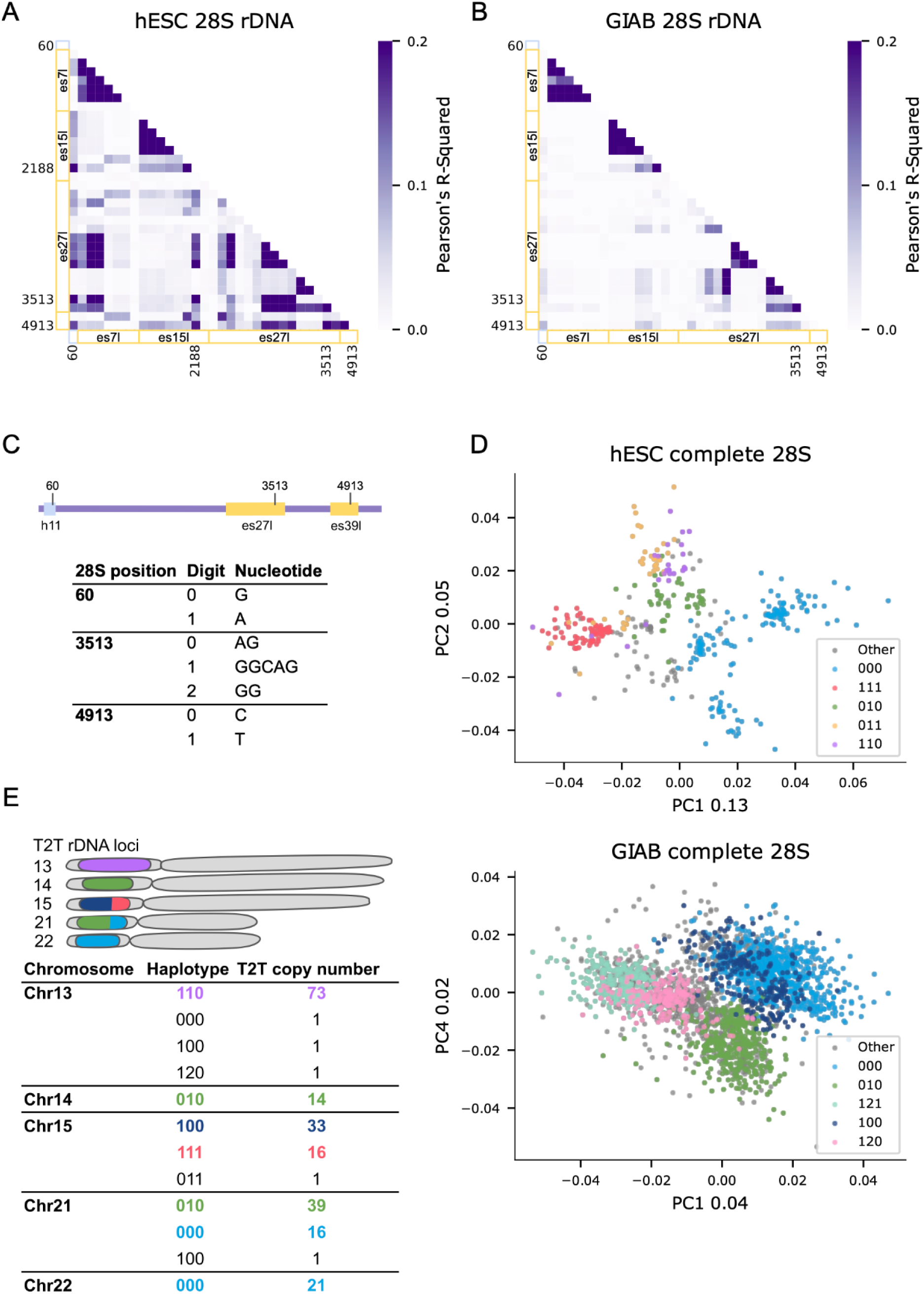
28S subtypes found by haplotype analysis. A. Correlation coefficient (Pearson’s r^2^) heatmap between positions across H7-hESC 28S rDNA with variant frequency >10%. X-axis and Y-axis are annotated by regions. Helix regions are annotated by light blue and ES regions are annotated by yellow. Individual positions with higher r^2^ between regions are also indicated. B. Same as (a) for the Genome In A Bottle (GIAB) dataset. C. Haplotype digit code to variant sequence conversion at the 3 positions with higher r^2^ in (a) and (b) D. Bray-Curtis Principal Coordinate Analysis (PCoA) of 386 H7-hESC 28S rDNA sequences (upper panel) and 386 28S rDNA sequences from each GIAB sample (lower panel). Each dot is a complete 28S rDNA sequence with similarity between sequences measured on 6-mers. The colors correspond to coloring an rDNA sequence by its 3 position haplotype described in (c). Numbers in the X and Y labels represent the PCoA explained variance. E. Telomere-to-telomere (T2T) haplotype distribution across the 5 acrocentric chromosomes. The rDNA acrocentric arms are presented in a schematic cartoon with proportions of rDNA haplotypes in different colors as found in the matching table below. Haplotypes match the 3 position haplotypes in (c). We indicate the rDNA copy number of each haplotype in every chromosome.

Previously it was shown that the rDNA array is composed of highly homogenized tandem clusters ^51^. We therefore next asked whether different 28S haplotypes are spatially separated in the genome as different subtypes. For the H7-hESC we have 386 complete 28S rDNA sequences and in the GIAB dataset we randomly subsampled each GIAB sample to 386 complete 28S rDNA sequences. For these datasets, we do not know rDNA-chromosome positioning. Notably, by comparison of 28S rDNA sequence similarities, we detected distinct 28S sequence groups in both hESC and GIAB (**Figure 3D**, Principal Coordinate Analysis, PCoA, of Bray-Curtis dissimilarities between 28S sequences ^52^, **Methods**). Here, the different clusters in PCoA space match different 28S haplotypes. Specifically, we observed that 28S sequences of a given haplotype are more similar to one another in their entire sequence compared to 28S rRNAs of other haplotypes (**Figure 3D**). When plotting individual haplotypes, there is less observed structure in individual haplotypes as compared to the combined data (**Figure S15**). In the T2T assembly, rDNA copies have chromosome coordinates, which enables us to measure 28S subtype presence at the five acrocentric chromosomes. Remarkably, we discovered that 28S haplotypes are largely chromosome specific (**Figure 3E**). When analyzing the 1KGP, we find that all of the haplotypes found in high frequency in the H7-hESC, GIAB and the T2T CHM13 genome, are also present in the 1KGP genomes. However, when examining haplotype frequency changes, we find high standard deviation between their frequencies across individuals (**Table S18**). This variability limits our understanding of the 28S haplotypes, which may be caused by high rates of nonallelic gene conversion across rDNA copies. Taken together, our result support that 28S haplotypes are genomically separated and belong to different subtypes.

### Ribosomes of different 28S subtypes have different structures

We next asked if different 28S subtypes have different ribosome structures. Notably, the abundant variants in the hESC were found in four different ES regions which were never previously resolved by CryoEM. Here, we treated the hESC sample with DMS, which covalently modifies the RNA at regions where the rRNA is accessible to allow for structure probing of the RNA (**Methods**). Using our RIBO-RT method for sequencing full length 28S with our RGA variant calling on DMS treated hESC cells, we obtained an accessibility map of the 28S. Importantly, we are able to predict the structure of the full length ES regions which was not previously possible.

We compared the two most abundant 28S subtypes and their linked variants and found they have different structures (**Figure 4A-C**, with 22% and 30% frequency of subtypes 1 and 2 respectively, **Methods**). While our method with DMS results in a full length accessibility map of the 28S, secondary structure prediction becomes less accurate for long RNA sequences. Given that ESs have tentacle-like extensions that protrude from the ribosome, we assumed that the core non-ES rRNA is not affected by changes in the ES regions which allowed us to focus on the structures of individual ESs. Most interestingly, we discovered that the ESs es7l, es15l and es27l have major DMS accessibility and structure differences when comparing the two subtypes observed at the GGC sites in es7l, es15l and es27l (**Figure 4A**, ES region box annotations, **Figure S16-20** for es7l, es15l and es27l). When focusing on es27l, the second longest ES, we noticed that the largest accessibility difference between the subtypes was at the site where es27l subtypes differ, at the GGC indel. Specifically, the subtype with one fewer tandem-repeat GGC insertion before the AG at position 3513 of the 28S showed greater DMS accessibility at position 3513 and its vicinity (**Figure 4D,E**). This GGC expands a six tandem-repeat GGCs, i.e. (GGC)_6_, which changes the region’s structure. Moreover, for the es27l region we found local structure changes near the sequence variants which opens the possibility that there are proteins or transcripts that may interact with the subtype with the GGCAG variant but not with the AG variant (**Figure 4D,E** region marked in red). Taken together, our DMS results provide evidence of structural differences for different ribosome subtypes.

**Figure 4.**
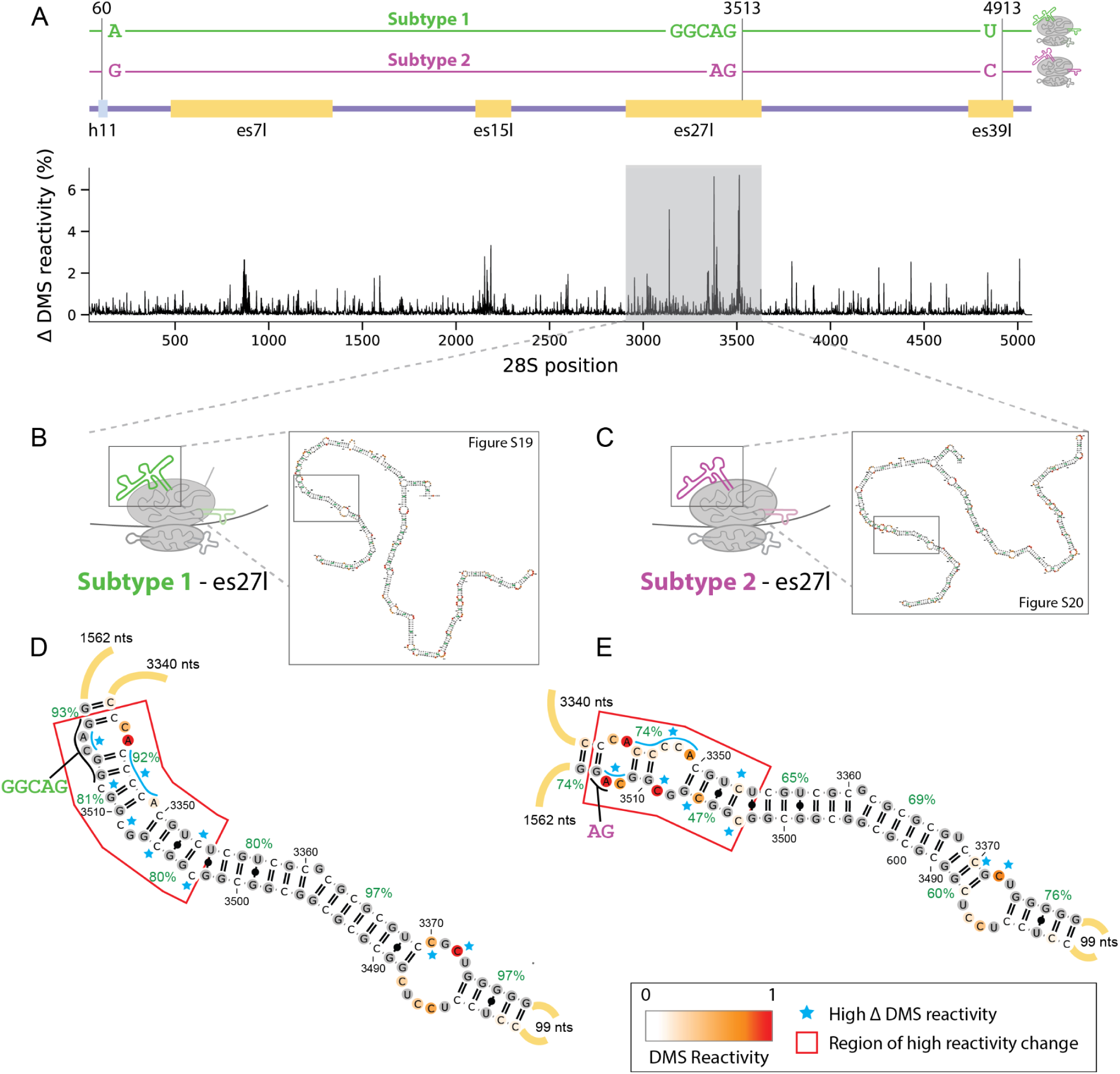
In-cell dimethyl sulfate (DMS) with long-read sequencing shows that 28S subtypes have different RNA 2D structure. A. Changes in DMS accessibility between two most abundant 28S subtypes across the complete 28S molecule. Subtypes are defined by the sequence variants observed at position 60 (28S:h11), 3513 (es27l), and 4913 (es39l), according to the numbering in NR_146117.1. Above is an illustration of the two subtypes, together with the annotations for the aforementioned regions and other regions with large differences in accessibility. X-axis is the nucleotide position along the 28S. Y-axis is the absolute percentage of DMS accessibility differences at a given position for a window size of 10 nucleotides. B. Illustration of es27l predicted secondary structure for subtype 1 (A, GGCAG, T). Detailed RNA 2D structure of the whole subtype 1 es27l is shown in Figure S19. C. Illustration of es27l predicted secondary structure for subtype 2 (G, AG, C). Detailed RNA 2D structure of the whole subtype 2 es27l is shown in Figure S20. D. Zoomed-in predicted RNA secondary structure of subtype 1 - es27l between position 3310 to 3552(+3). RNA secondary structures are colored by DMS reactivity and helix confidence estimates are depicted as green percentages. Regions with major differences are annotated by the red box. Nucleotides with differing accessibility between the two subtypes are highlighted by blue stars. E. Zoomed-in predicted RNA secondary structure of subtype 2 - es27l between position 3310 to 3552. Detailed description of annotation is the same as (d).

### Quantifying the relative abundance of rRNA variants in expression data

Previously, 10 rRNA variants were annotated and showed changed expression between mouse tissues^5^. Here, we found that one of these rRNA variants replicates in our atlas. This promped us to check if rRNA variations accross human tissues by analyzing the publicly available Genotype-Tissue Expression (GTEx) short-read RNA-seq dataset to test if rRNA variant frequencies are associated with human tissue biology (**Methods** for altas usage instructions). Previous studies comparing mRNA across tissues in the GTEx dataset found tissue-specific, including brain-specific, gene expression ^53,54^. Here we analyzed 2,618 samples from 332 individuals and 44 tissues from GTEx and asked if rRNA subtypes differ in their expression in these tissues (**Figure 5A**). We hypothesized that the rRNA subtypes that we identified as highly expressed in the hESC may be important for tissue development. Strikingly, the most abundant subtypes in GTEx significantly differed in expression in many tissues (**Figure 5B-D, upper panels, Figure S21-25**, and **Table S19-20**: P-value < 0.05 FDR corrected Mann-Whitney U rank sum test). Notably, when comparing subtypes expression levels across tissues, we observed significant differences between tissues derived from the ectoderm and endoderm germ layers (**Figure 5B-D, lower panels, Table S21** FDR corrected ranksums test comparing subtype relative abundances of ectoderm-derived tissues in blue and endoderm-derived tissues in red). Most of the ectoderm-derived tissues belong to brain tissues, and most endoderm derived tissues are digestive-system tissues (**Figure 5A** endoderm and ectoderm derived tissues are labeled). Taken together, our results support major changes in the expression of rRNA subtypes across tissues.

**Figure 5.**
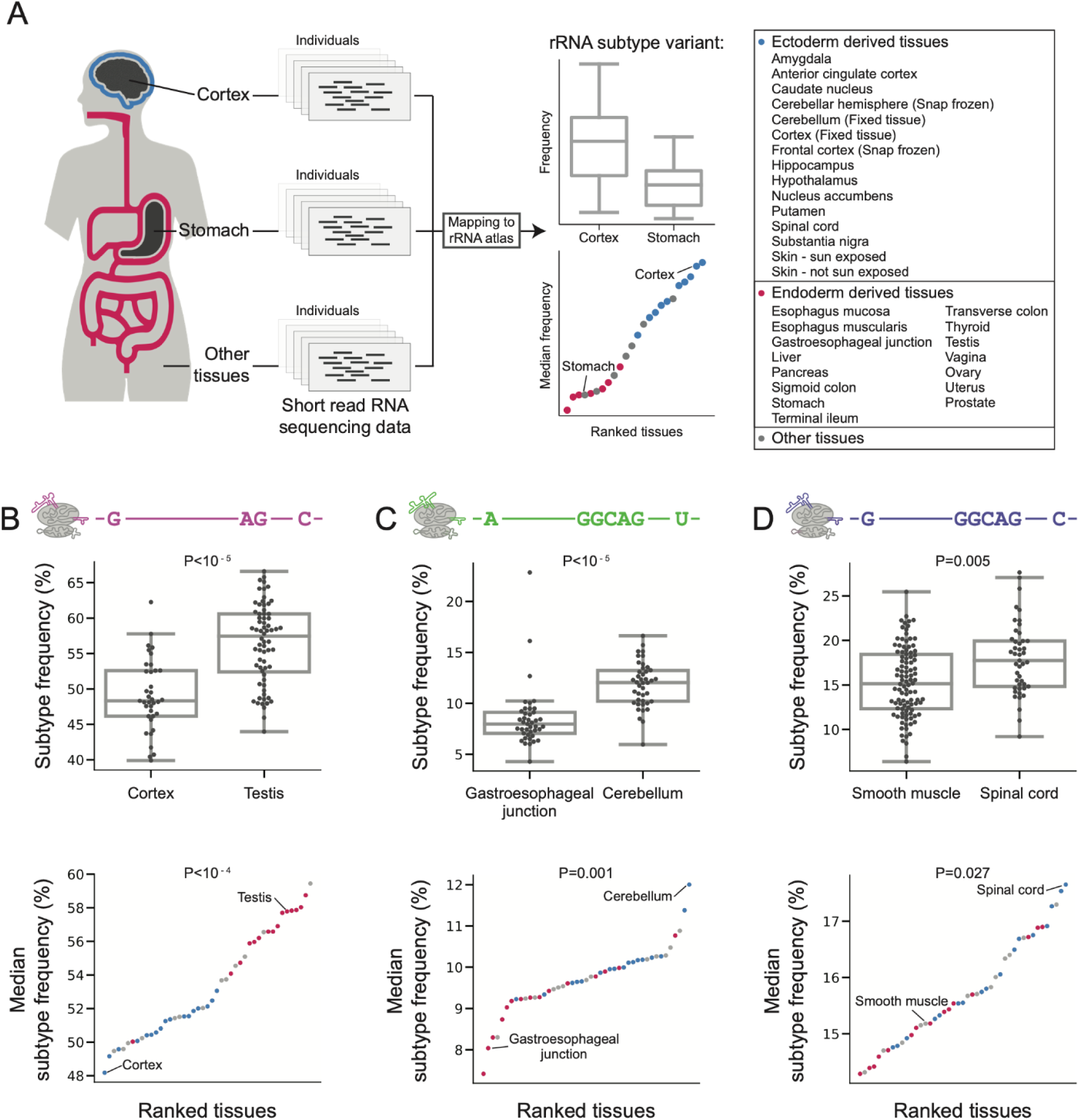
rRNA subtype expression is tissue specific and differs between tissues derived from the ectoderm and endoderm lineages. A. Schematic representation of the GTEx analysis focusing on Cortex versus Stomach comparison. We map rRNA reads from different samples to rRNA subtypes. Per rRNA subtype, we illustrate variant expression comparison between tissues: Upper panel with a box plot comparing rRNA subtype expression in Cortex and Stomach samples. Bottom panel shows the median rRNA subtype expression across all tissues. Cortex and Stomach are annotated, and all ectoderm and endoderm tissues are highlighted in blue/red marks. B. Upper panel: Box plot comparing the expression levels of the rRNA subtype with the haplotype G,AG,C (positions 60,3513,4913) in Cortex and Testis samples. Bottom panel: Scatter plot showing the median frequency of the rRNA subtype from the upper panel across all tissues. Tissues derived from ectoderm are marked in blue, tissues derived from endoderm are marked in red, other tissues in gray. The Cortex and Testis that were shown in the top panel are annotated with a line. C. Same as (b) for the rRNA subtype with the haplotype A,GGCAG,T highlighting Gastroesophageal Junction and Cerebellum samples. D. Same as (b) for the rRNA subtype with the haplotype G,GGCAG,C highlighting Smooth muscle and Spinal cord samples.

Lastly, we asked if changes in the expression of rRNA variants are associated with cancer. For this, we used 10,030 samples of short-read RNA-seq with clinical phenotypes from The Cancer Genome Atlas (TCGA) ^55^. When comparing cancer types, we found distinct expression patterns of rRNA regional variants across cancers (**Figure S26-31, Table S22** for region annotations). To test if rRNA variants are cancer-specific, we compared cancer biopsies to control biopsies from the same tissues. Surprisingly, we identified specific rRNA regional variants with significantly different expression levels in control and in cancer biopsies for 11 cancer types (**Figure 6**, **Table S22** for alternative allele regional variant abundances**, Table S23**, P-value < 0.05 after FDR correction, bootstrapping 10,000 times Mann-Whitney U rank sum test with sub sampling controls to match the number of cancer samples). These include rRNA variants that while they appeared in low abundance in both the H7-hESC and control biopsies, are found elevated in cancer biopsies. Thus even low abundance variants hold immense importance as disease biomarkers.

**Figure 6.**
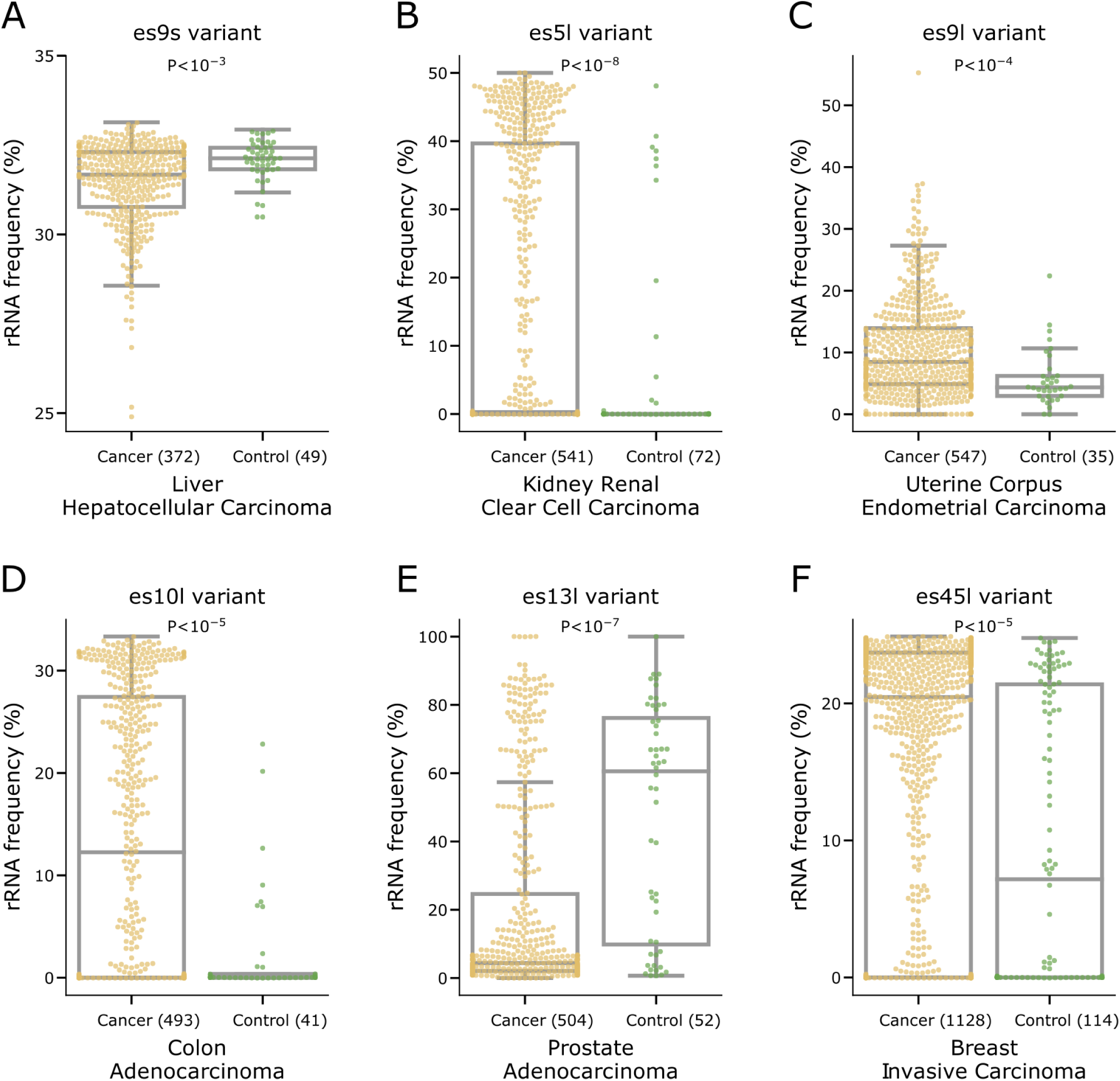
Cancer-specific rRNA variant expression. A. Box plot showing the distribution of rRNA frequencies of the top expressed, alternate allele regional variant, of es9s, ES:es9s:6_d14_r115, across TCGA cancer and control samples for Liver Hepatocellular Carcinoma. The box plot is overlaid with a categorical scatter plot which saturates at values with over 20 samples and not all points are displayed. The sample sizes of cancer and controls is indicated in brackets and the P-value is indicated at the top (**Table S23**). B. Same as (a) for region es5l, ES:es5l:12_d1_r1, in Kidney Renal Clear Cell Carcinoma. C. Same as (a) for region es9l, ES:es9l:21_d2_r8, in Uterine Corpus Endometrial Carcinoma. D. Same as (a) for region es10l, ES:es9l:21_d2_r8, in Colon Carcinoma. E. Same as (a) for region es13l, ES:es13l:2_d3_r141, in Prostate Adenocarcinoma. F. Same as (a) for region es45l, ES:es45l:7_d2_r45, in Breast Invasive Carcinoma.

We conclude that our atlas enables direct measuring of rRNA variants changes in expression data. Moreover, we showed that atlas variants are present in translating ribosomes and that they are differentially expressed across tissues and cancer types.

## Discussion

Here, by developing a pipeline for long-read sequencing and analysis of rDNA and rRNA from actively translating ribosomes, we measured for the first time variant frequencies in rDNA and rRNA and used *in-situ* sequencing microscopy to validate co-variant expression in individual cells. With this atlas we have enabled greater understanding of the often neglected yet ubiquitous rRNA sequencing data and built an atlas of functional human 18S and 28S rRNA variants at different resolutions, from nucleotide position variants, to 28S gene-level subtypes as a useful resource for studying rRNA variations, and composition across biological conditions (**Extended data 1-5**).

In our study, we have discovered chromosome-associated rDNA subtypes, revealing that different ribosome subtypes based on rRNA sequence variation exist. It may be possible that spatial separation of rDNA subtypes enable regulation of their expression at the chromosome level through allelic inactivation of rDNA loci or inactivation of nucleolar organizer regions (NORs) in the distal junction ^56–58^. This may enable global remodeling of rDNA transcription and promote specific ribosome subtypes to be expressed within individual cells. Additionally, using DMS structure probing of full length 28S, we discovered that different rRNA subtypes have different structures at ES regions including different DMS accessibility profiles. Since these ES regions are solvent-exposed and highly flexible, these ES variations may fine-tune regulation of mRNA translation based on differential association with ribosome-associated proteins, mRNA transcripts, or other factors. Moreover, by analyzing the GTEx dataset, we observed differential expression of rRNA subtypes between tissues belonging to ectoderm and endoderm lineages. This pattern might hint at specialized functions of different ribosome subtypes. Long-lived cells associated with the nervous system might require ribosome subtypes that emphasize translation fidelity over speed, as compared to rapidly dividing cells in the digestive tract that require constant replacement given harsh local environments. Indeed, our lab and others have previously shown that es27l plays a role in translation fidelity through association with ribosome-associated proteins ^44,59,60^. Such interactions that trade speed over fidelity might be fine-tuned by the expression of different rRNA subtypes.

Finally, by analyzing the TCGA dataset we discovered that some low abundant rRNA variants in control biopsies were elevated in cancer biopsies. Yet the mechanism of elevated expression of such variations remains unknown. One possible mechanism may be enhanced transcription of specific rDNA copies bearing coding sequence variants and interestingly it was shown that Lung Adenocarcinoma samples were enriched with somatic and germline mutations at rDNA promoter regions ^61^. Alternatively, de novo somatic mutations may increase certain rDNA variant frequencies. Future work is needed to understand whether they promote oncogenic ribosome activity and how they are regulated. Therefore, our results provide another layer of ribosome specificity wherein cancer cells might deploy a particular rRNA variant that is more compatible with their cellular fitness. Importantly, we found that specific rRNA variants may be used as biomarkers for disease. Notably, 5-fluorouracil, a common chemotherapy drug, was recently shown to incorporate into rRNA and promote drug resistance by changing mRNA translation ^62^. It may be that drugs directly target specific rRNA variants and further examination would be needed to test whether they should be used for cancer specific therapies. Together, our results reveal the presence of structurally different ribosomes at the level of rRNA and provide the first atlas to distinguish different types of ribosomes and to link them to different cellular programs, including those underlying human health and disease.

## Supporting information

Complete Atlas

## Acknowledgments

We thank Rhiju Das for interpreting DMS results. We thank Xiangling Meng, Craig Kerr, Ali Wilkening and Michael Montgomery for help with early experiments that were not later followed in this study. We thank the Barna and Pritchard group members for discussions.

MB is supported by the New York Stem Cell Foundation and the National Institutes of Health grant R01HD086634. JKP is supported by RO1 HG008140. DR is supported by ALTF 1042-2019 EMBO and LT000218/2020-L HFSP postdoctoral scholarships. TTS is supported by a National Science Scholarship (PhD) from the Agency for Science, Technology and Research. MB is a NYSCF Robertson Investigator. XW is supported by Edward Scolnick Professorship, Ono Pharma Breakthrough Science Initiative Award, Merkin Institute Fellowship, and NIH DP2 New Innovator Award 1DP2GM146245-01.

## Authors contributions

DR conceived the project, designed experimental and computational analyzes, conducted all computational analyses, interpreted the results, and wrote the manuscript. TTS designed and conducted all polysome and DMS sequencing experiments, interpreted the results, and wrote the manuscript. XS developed, optimized, and performed SWITCH-seq experiments. XW supervised the in-situ sequencing experiment. JPS helped in computational analyses. NG helped in experimental data collection. NSA helped interpret GTEx data. RR helped in DMS analyses. MB and JKP conceived and directed the project and analyses, designed the analyses, interpreted the results and wrote the manuscript.

## Declaration of interests

XW is a scientific cofounder of Stellaromics. X.W. and X.S. are inventors on pending patent applications related to SWITCH-seq

## Data and materials availability

The atlas is available as Extended Data to this publication.

H7-hESC raw rDNA and rRNA sequencing data is available under BioProject ID PRJNA926787.

## Limitations of study

In this paper we created an atlas of human rRNA sequence variations in translating ribosomes which we correlate with both development as well as to cancer. In this study we do not demonstrate that expression differences of rRNA variants have functional implications on human development and disease. In the TCGA dataset control samples do not belong to the same matched cancer biopsy and some cancer types have low control sample sizes. In our haplotype analysis we found high standard deviation between haplotype frequencies across individuals which limits our understanding of their functional significance. The expressed rRNA variants belong to the H7-hESC and K562 cell lines. It is likely that there are rRNA variants that are expressed in other human cells or samples that are not found in these cell lines. The RGA method is not limited to variant discovery between paralog genes, it can be applied for variant discovery between any related sequences for example in detecting variants between amplicon sequences.

## Notes

### Summary of Updates

Two more analyses were added to the manuscript.

