## Supplementary figures and methods for "Diversity of ribosomes at the level of rRNA variation associated with human health and disease"

### s24Supplementary Figures

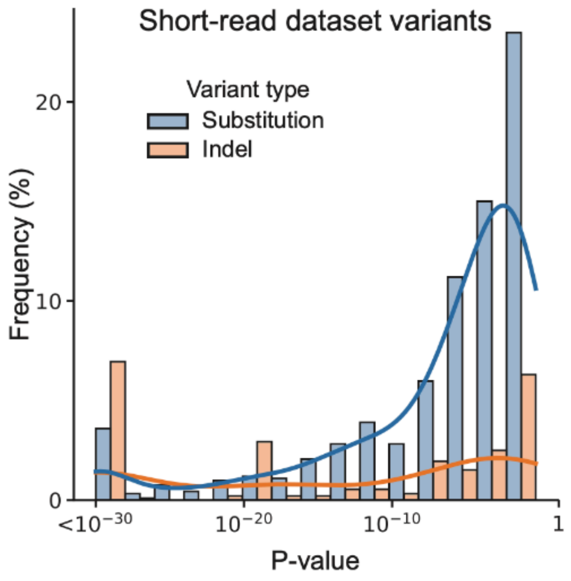


**Figure S1. Indels are associated with lower P-values compared to SNVs**

Distribution of Mutect2 false discovery rate (FDR)-corrected log 10 likelihood ratio scores of variant existence measured for substitutions and indels.

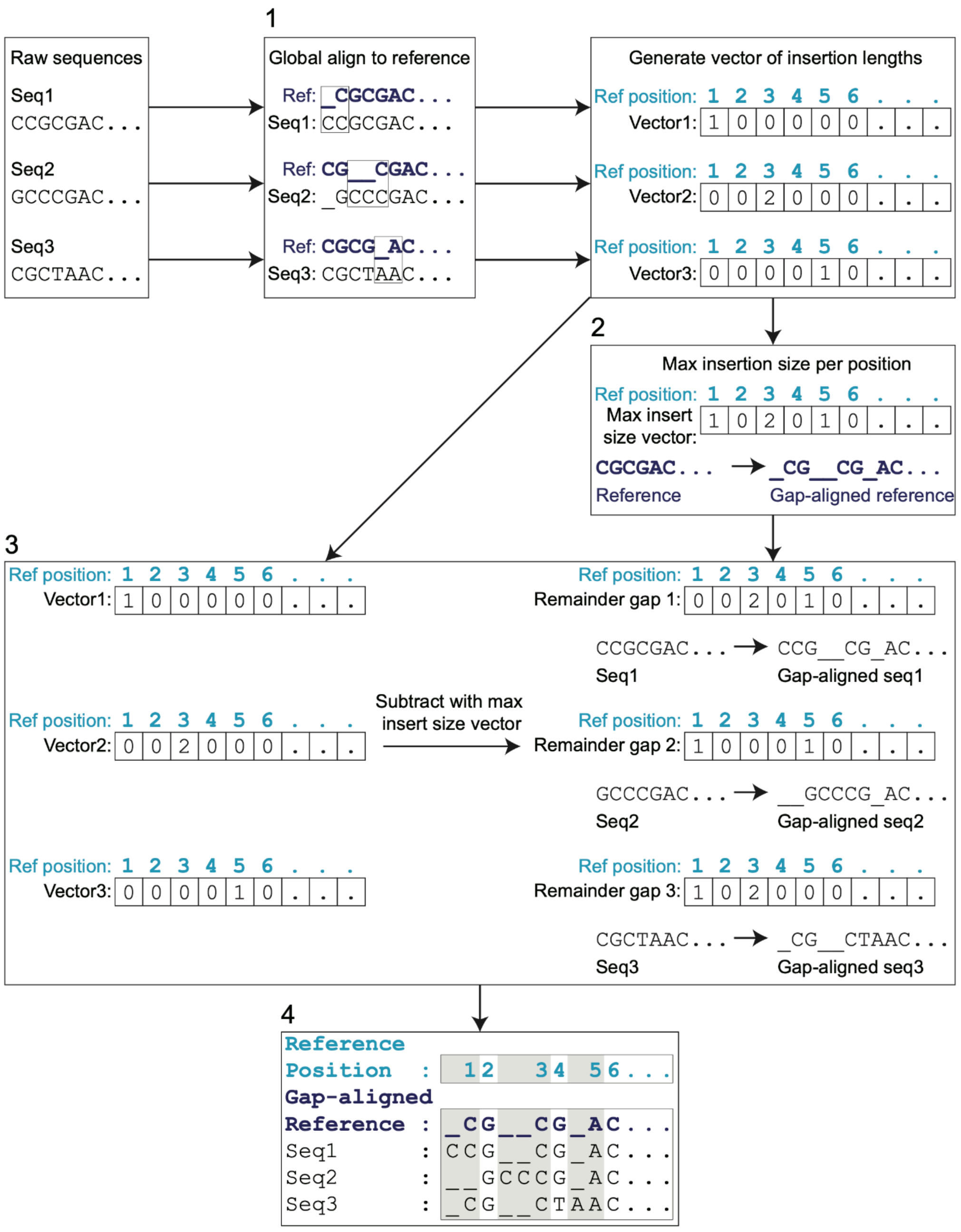


**Figure S2. Reference Gap Alignment (RGA) steps**

The four steps in the RGA method are illustrated with matching numbers to the steps in the main text.

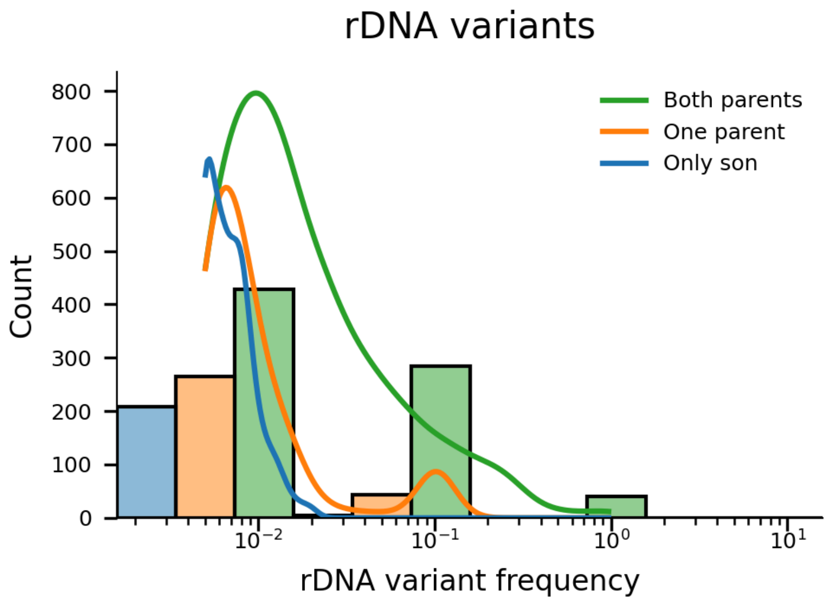


**Figure S3. rDNA variants frequecies from a progeny sample support variant heritability**

rDNA variant frequencies of the GIAB Ashkenazy son are shown in a histogram and are color coded in blue if the are not found in neither parents, orange if found in only one of the parents and in green if found in both.

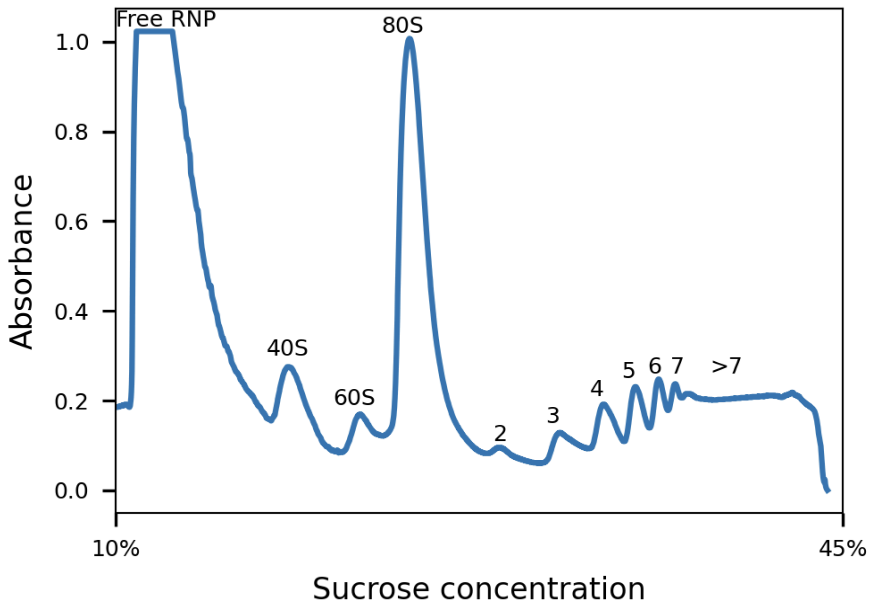

**Figure S4. polysome profile from H7-hESC 10-45% sucrose gradient fractionation**

H7-hESC A260 trace showing the free ribonucleoproteins (RNP), free 40S and 60S subunits, 80S monosomes, and polysomes (marked with 2-7 and >7).

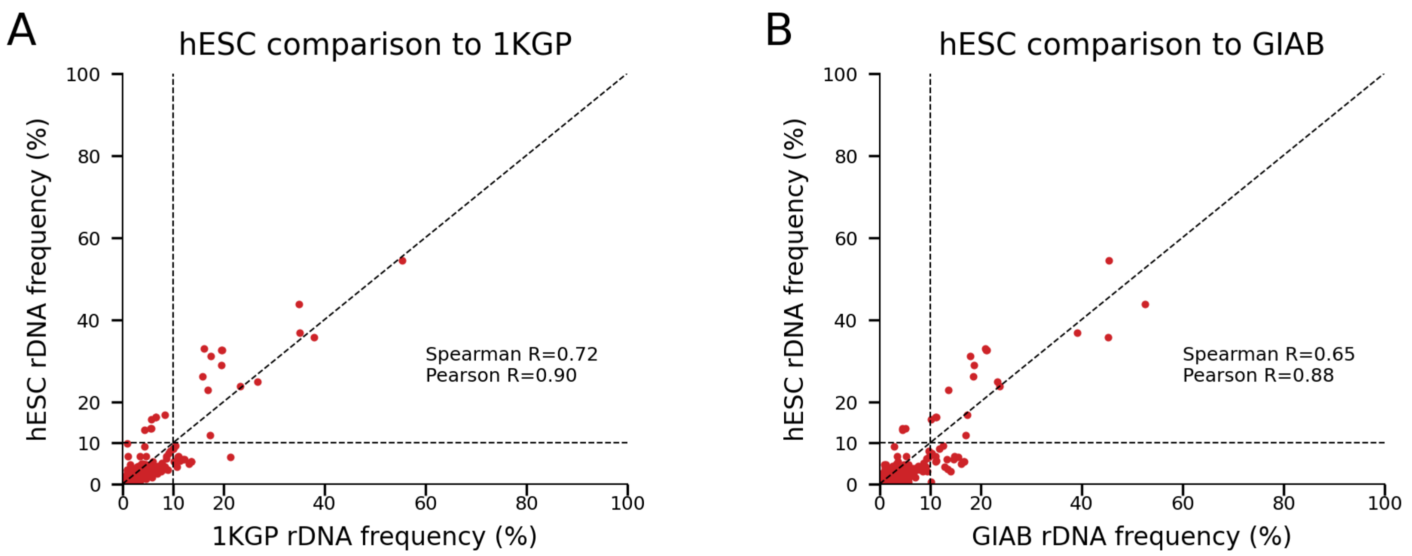


**Figure S5: 28S rDNA variant frequencies comparing 1KGP and GIAB to the H7-hESC**

A. Scatter plot comparing 28S rDNA variant frequencies found in long-read of the 1KGP (x-axis) and in the H7-hESC (y-axis). Pearson and Spearman correlations are indicated.
B. Same as (A) for GIAB (x-axis).

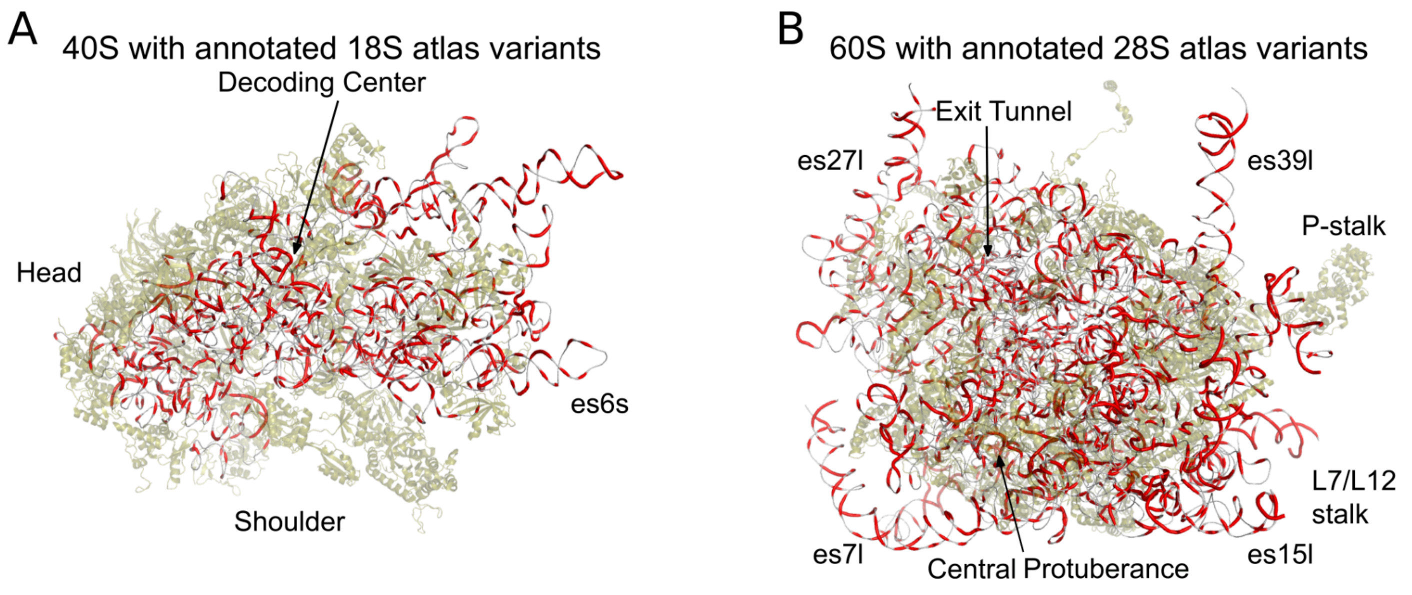

**Figure S6. 3D structure of the 40S and 60S with variants**

A. 40S 3D structure with 18S atlas variants. Ribosomal proteins are presented in semitransparent green, 18S rRNA in gray. Positions with variants are highlighted in red.
B. Same as (A) for the 60S and the 28S atlas variants.

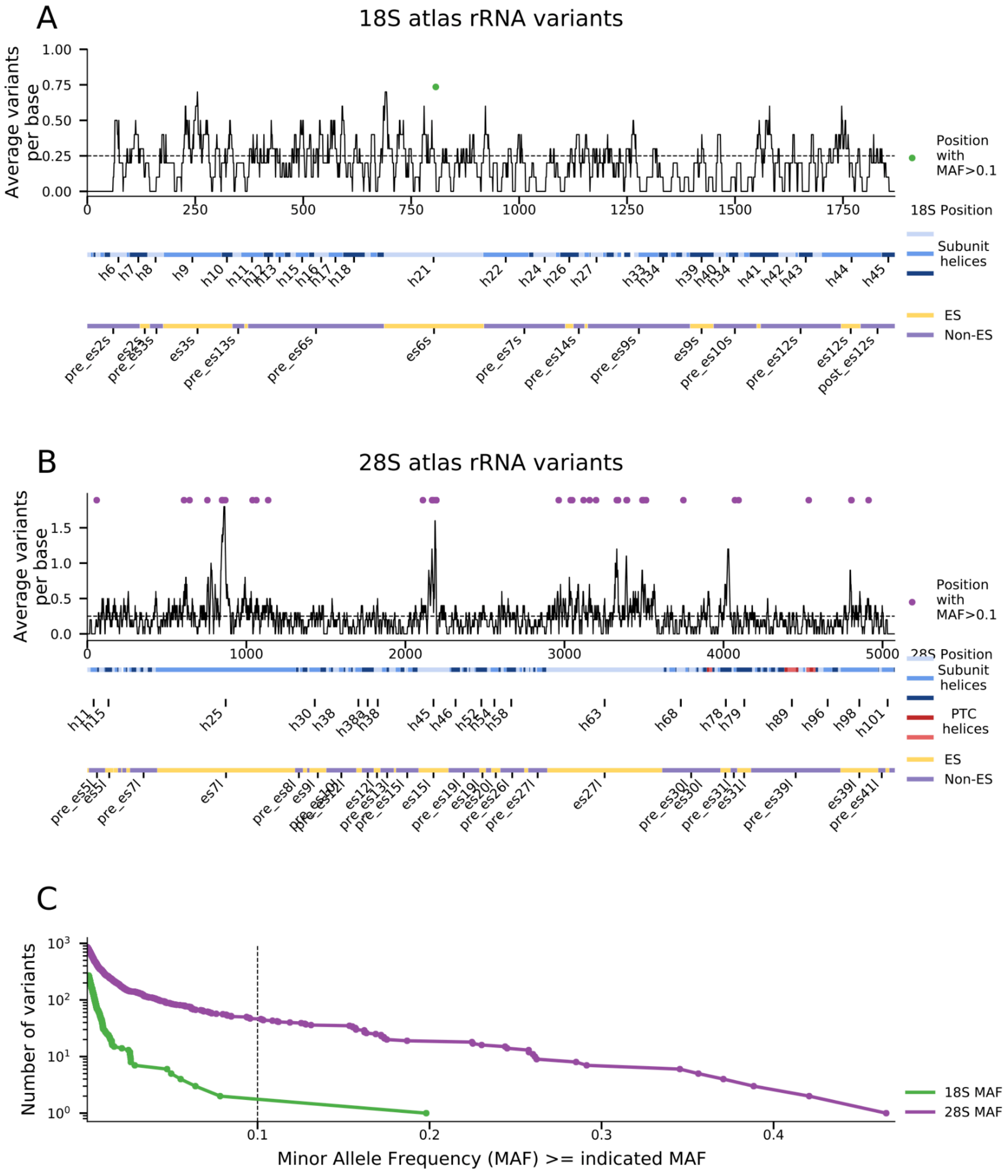


**Figure S7. 18S and 28S variant distributions**

A. Number of variants at 18S positions. X axis is the nucleotide position along the 18S. Below the X axis are annotations for 18S helix regions and ES regions. Y axis is the average number of atlas variants at a given position for a window size of 20 bases. Green dot at x = 807 (at helix h21 or es6s) is a position with a rRNA variant with minor allele frequency (MAF) >0.1. This dot is also plotted in panel (C).
B. Same as (A) but for the 28S. Purple dots annotate all positions with a variant with MAF>0.1. All purple dots are presented in panel (C).
C. rRNA allele frequency spectrum plot. Values in the X-axis indicate a MAF, and the Y axis are the number of variants with at least the X-axis matched MAF. For example, there are two variants with MAF >0.4 and 6 variants with MAF>0.3. The green plot matches 18S variants and purple plot matches the 28S variants. Individual variants are marked with a dot on the line plot. Dashed line indicates MAF equal to 0.1. The green and purple dots which are in panels (A) and (B) are on the right side of the dashed black line marking MAF=0.1.

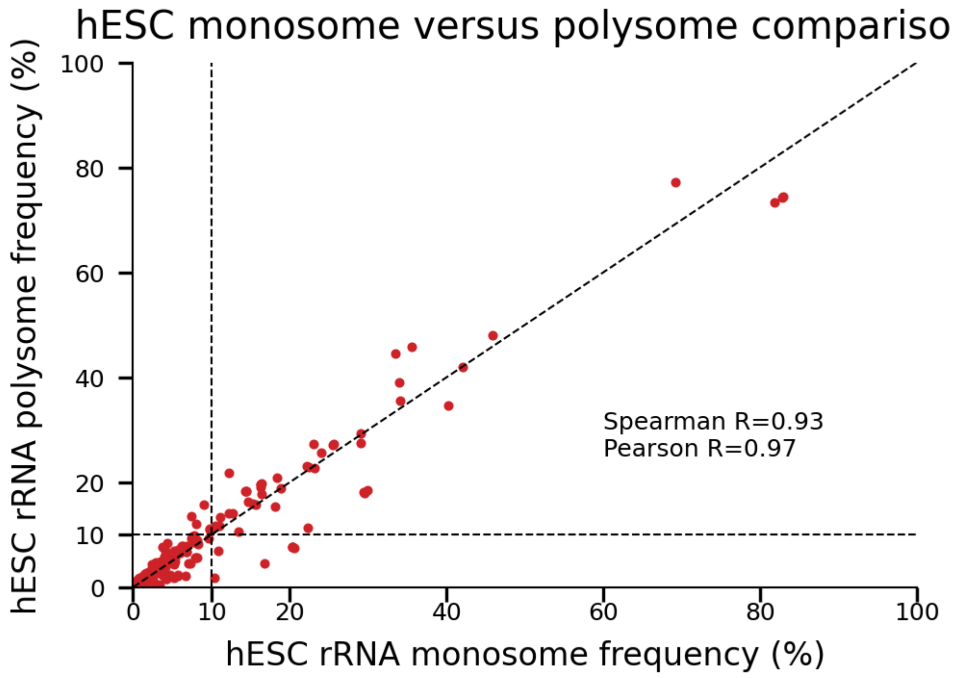


**Figure S8. H7-hESC 28S rRNA variant frequecies in monosomes and polysomes**

Scatter plot of 28S rRNA variant frequecies from the H7-hESC comparing polysome fractions. In the x-axis rRNA variant frequencies are calculared from monosomes and in the y-axis from polysomes. Pearson and Spearman correlations are indicated.

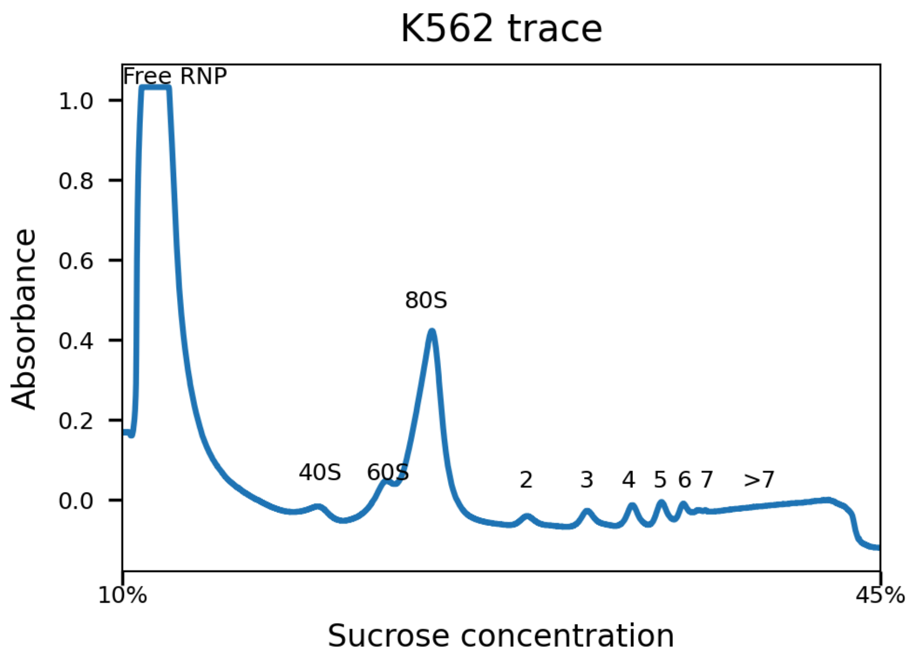


**Figure S9. polysome profile from K562 cell-line in 10-45% sucrose gradient fractionation**

K562 cancer cell-line A260 trace showing the free ribonucleoproteins (RNP), free 40S and 60S subunits, 80S monosomes, and polysomes (marked with 2-7 and >7).

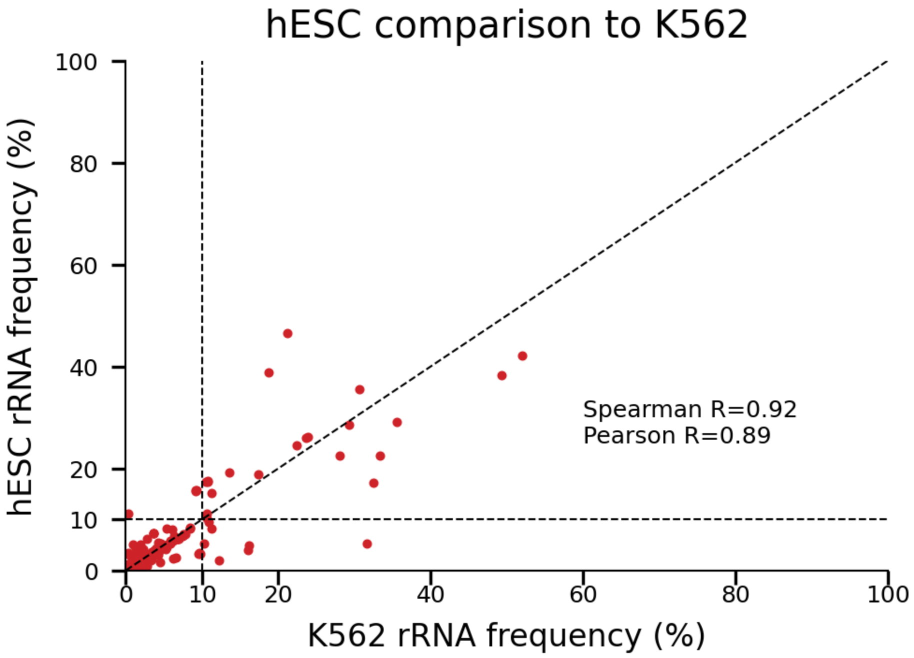

**Figure S10. 28S rRNA variant frequencies comparing K562 to the H7-hESC**

Scatter plot comparing 28S rRNA variant frequencies found in the K562 cancer cell-line (x-axis) and in the H7-hESC (y-axis). Pearson and Spearman correlations are indicated.

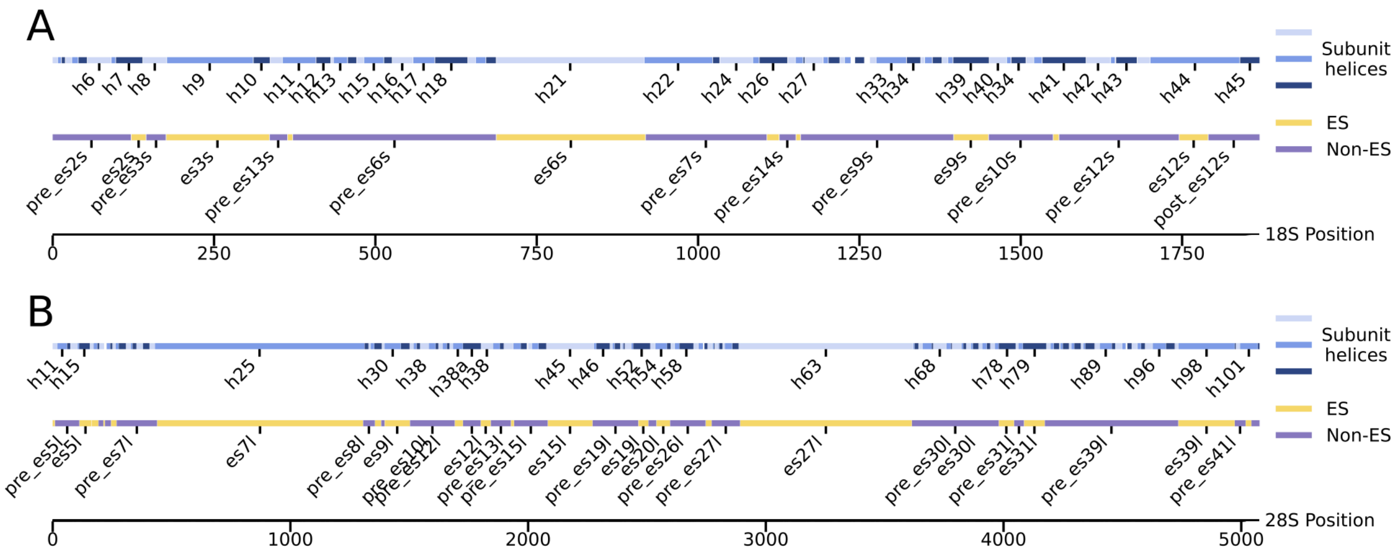


**Figure S11. 18S and 28S region annotation**

A. 18S helix and ES region annotation. Only helices and ES regions with at least 20 bases are labeled

B. 28S helix and ES region annotation. Only helices and ES regions with at least 40 bases are labeled

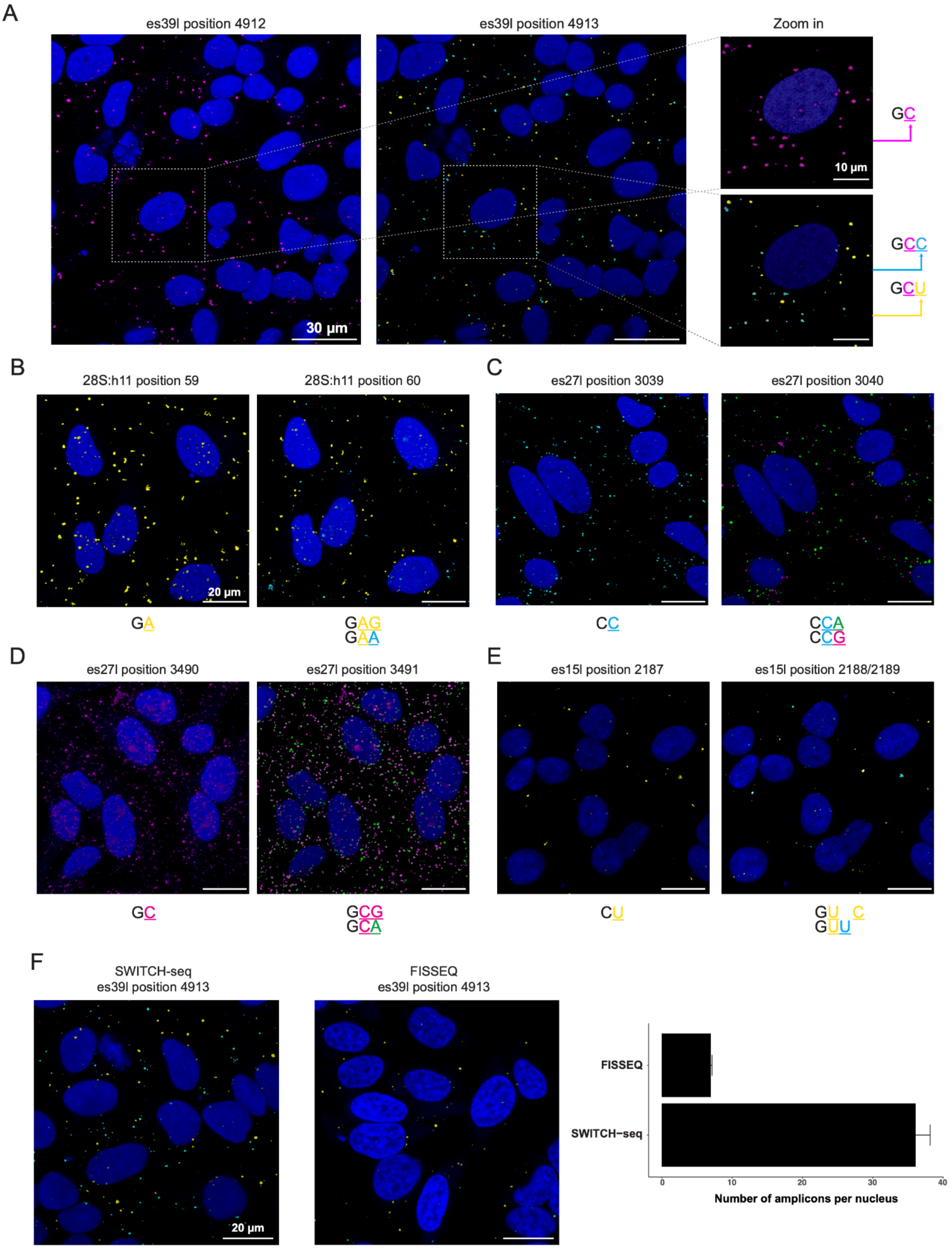
A
es39l position 4912
es39l position 4913
Zoom in
30 µm
10 µm
GC
GCC
GCU
B
28S:h11 position 59
28S:h11 position 60
20 µm
GA
GAG
GAA
C
es27l position 3039
es27l position 3040
CC
CCA
CCG
D
es27l position 3490
es27l position 3491
GC
GCG
GCA
E
es15l position 2187
es15l position 2188/2189
CU
GU C
GUU
F
SWITCH-seq
es39l position 4913
FISSEQ
es39l position 4913
20 µm
FISSEQ
SWITCH−seq
0
10
20
30
40
Number of amplicons per nucleus

**Figure S12. *In-situ* sequencing of rRNA variants**

A. Two rounds of representative fluorescent *in situ* sequencing images of HeLa cells (DAPI staining in blue) are presented for the es39l-probed region. We identified a non-variable base C (magenta) at position 4912. At position 4913, two alternative sequences were revealed: the known reference sequence C (cyan) and the alternative variant U (yellow).
B. Similar to (A) for 28S:h11 where G and A are detected at position 60
C. Similar to (A) for es27l where A and G are detected at position 3040
D. Similar to (A) for es27l where G and A are detected at position 3491
E. Similar to (A) for es15 where a U insertion is detected at position 2188.
F. Representative fluorescence images comparing SWITCH-seq and FISSEQ in the detection of the rRNA variant at es39l position 4913.

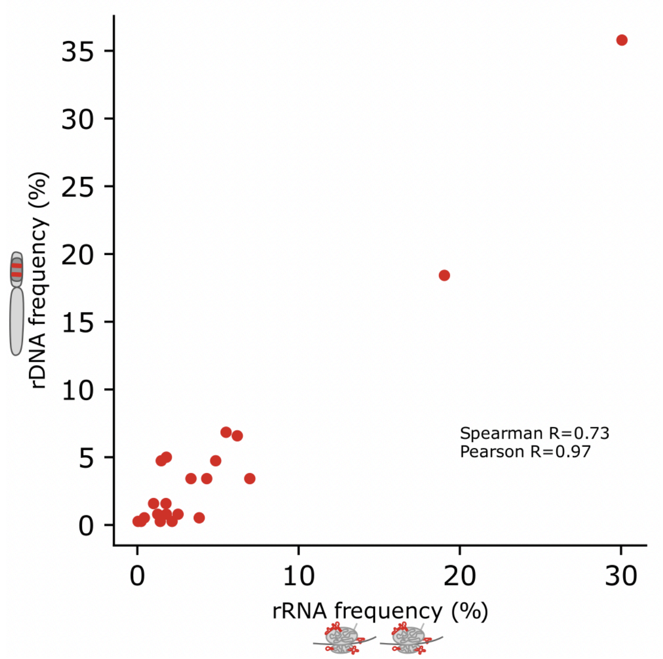


**Figure S13. 28S rRNA subtype frequencies in h7-hESC**

Scatter plot showing the frequencies of 28S rRNA subtypes in rRNA (x-axis) and rDNA (y-axis) in H7-hESC

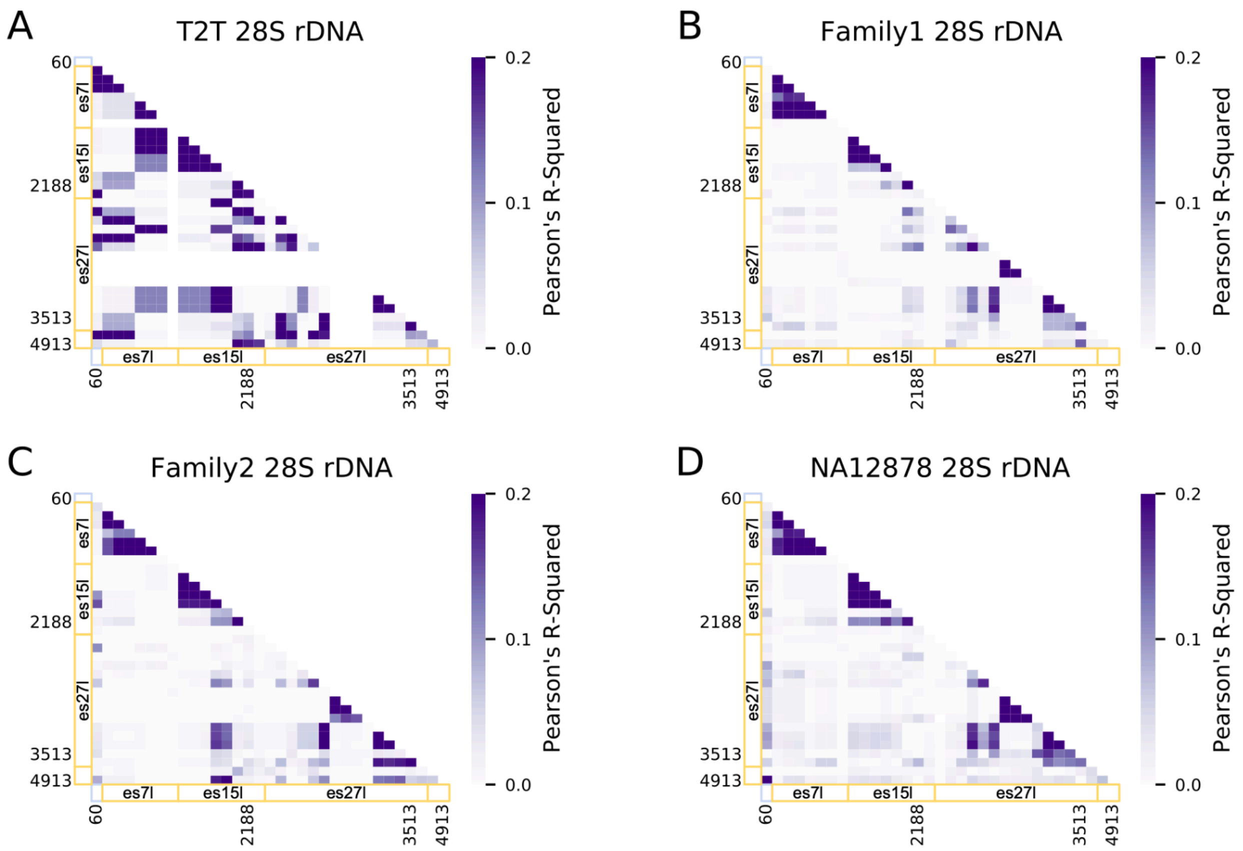


**Figure S14. 28S subtypes in T2T and GIAB**

A. Correlation coefficient (Pearson's $r^2$) heatmap of T2T 28S rDNA for the same positions analyzed in the h7-hESC (**Figure 1F, 3A**). X-axis and Y-axis are annotated by regions. Helix regions are annotated by light blue and ES regions are annotated by yellow. H7-hESC positions with higher $r^2$ are annotated.
B. Same as (A) for the Chinese Han family trio from the GIAB dataset.
C. Same as (A) for the Ashkenazi family trio from the GIAB dataset.
D. Same as (A) for the HA12878 cell line from the GIAB dataset.

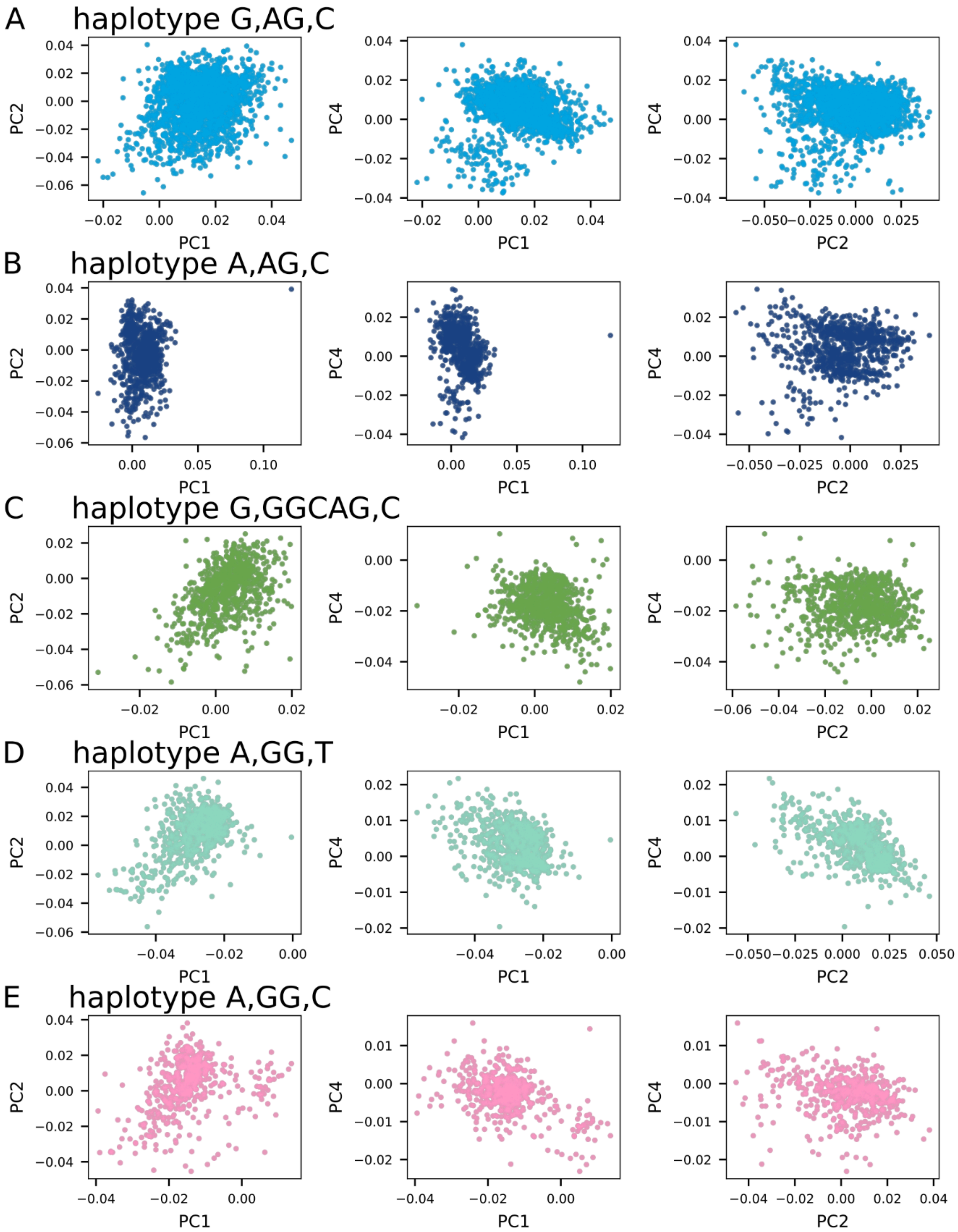
A
haplotype G,AG,C
B
haplotype A,AG,C
C
haplotype G,GGCAG,C
D
haplotype A,GG,T
E
haplotype A,GG,C
PC1
PC2
PC4

**Figure S15. 28S haplotype groups as shown by Bray-Curtis Principal Coordinate Analysis (PCoA)**

PCoA of 386 28S rDNA sequences from each GIAB sample. The first, second and fourth PCs are presented for 28S sequences that belong to different haplotypes where each haplotype is presented in a separte panel A-E. Each dot is a complete 28S rDNA sequence with similarity between sequences measured on 6-mers. The colors correspond to coloring an rDNA sequence by its 3 position haplotype shown in the main Figure 3D.

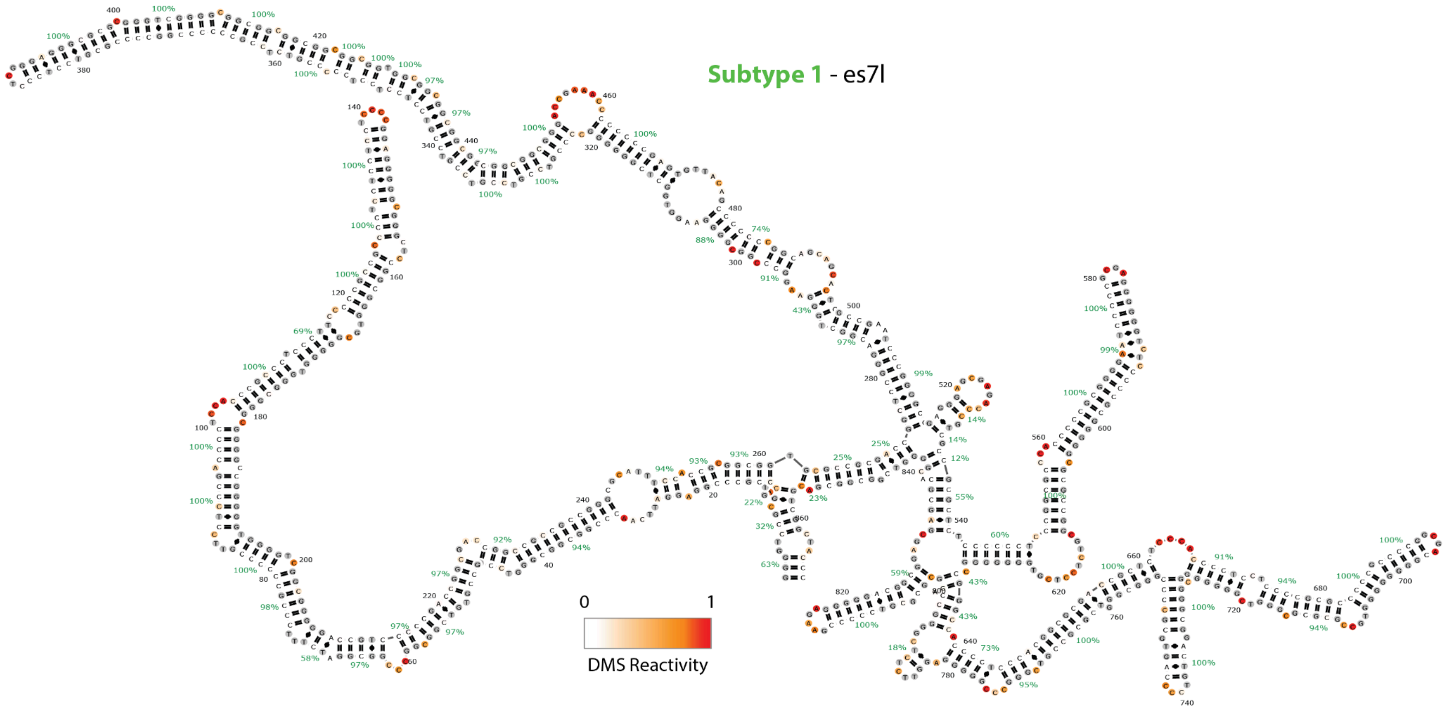

**Figure S16. In-cell DMS with long-read sequencing shows accessibility differences in es7l of different 28S subtypes**

RNA secondary structure of es7l predicted secondary structure for subtype 1 (A, GGCAG, T).

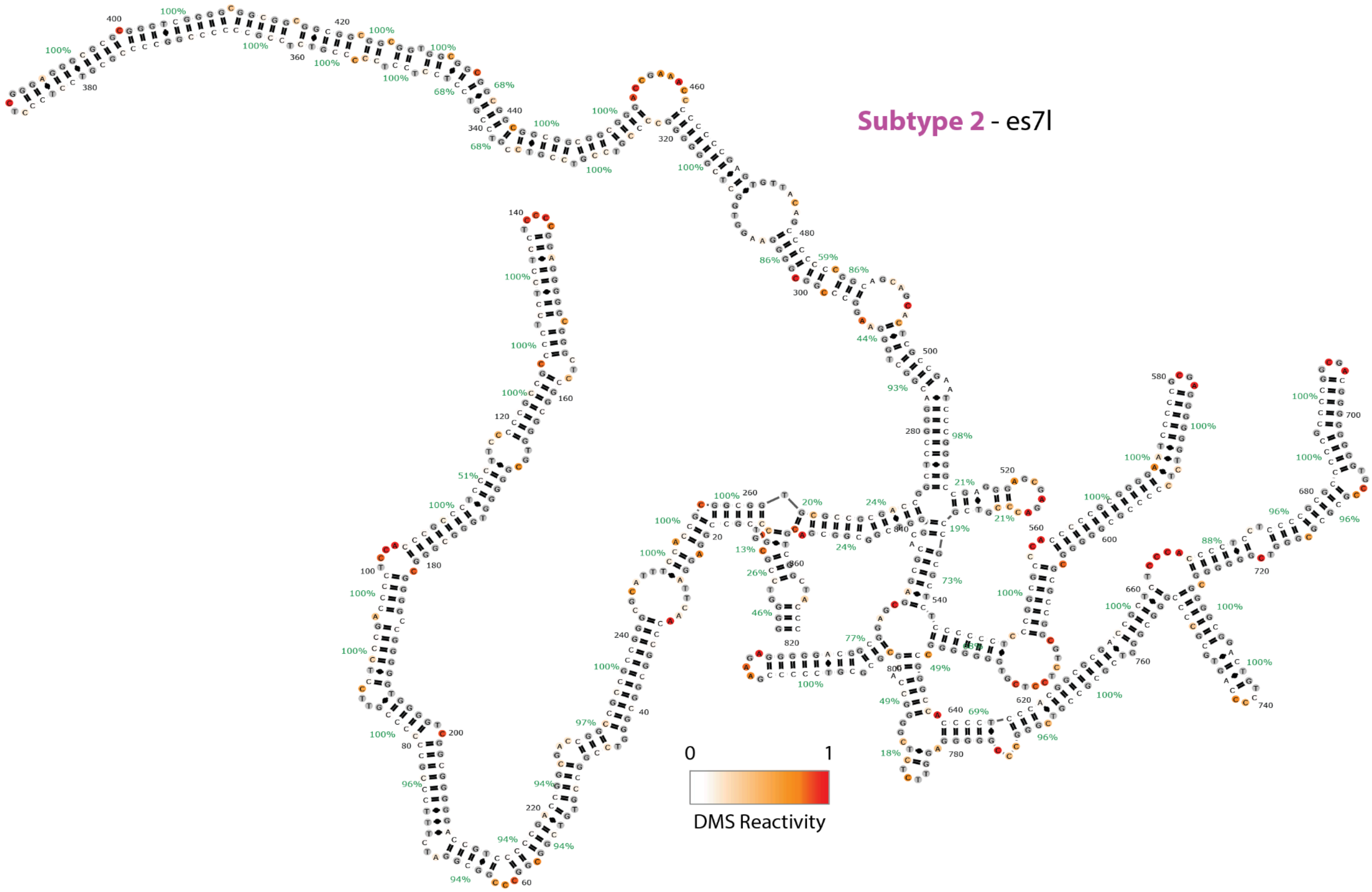


**Figure S17. In-cell DMS with long-read sequencing shows accessibility differences in es7l of different 28S subtypes**

RNA secondary structure of es7l predicted secondary structure for subtype 2 (G, AG, C).

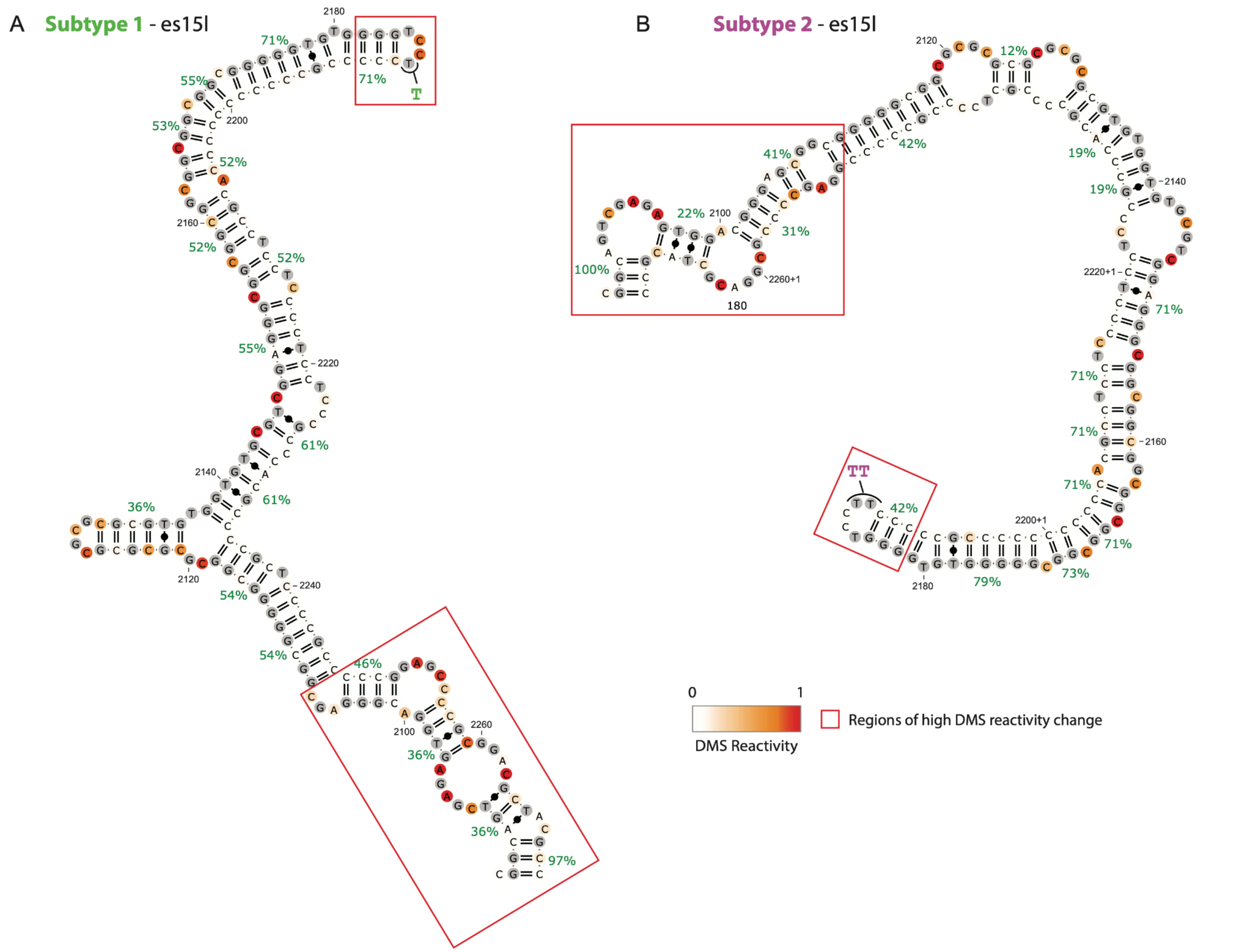


**Figure S18. In-cell DMS with long-read sequencing shows that es15l of different 28S subtypes have different RNA 2D structure**

A. RNA secondary structure of es15l predicted secondary structure for subtype 1 (A, GGCAG, T).
B. RNA secondary structure of es15l predicted secondary structure for subtype 2 (G, AG, C).

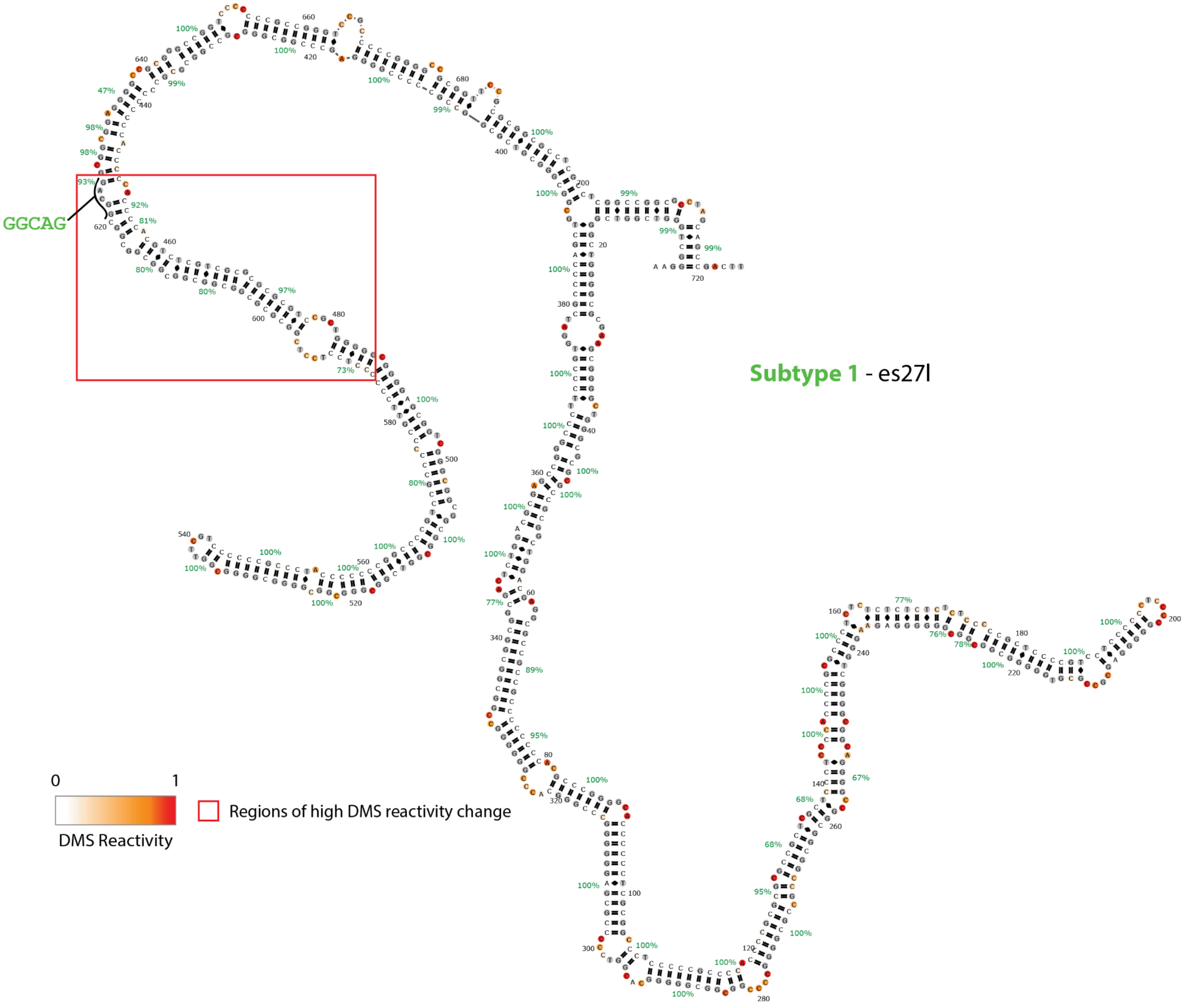


**Figure S19. In-cell DMS with long-read sequencing shows that es27l of different 28S subtypes have different RNA 2D structure**

RNA secondary structure of es15l predicted secondary structure for subtype 1 (A, GGCAG, T).

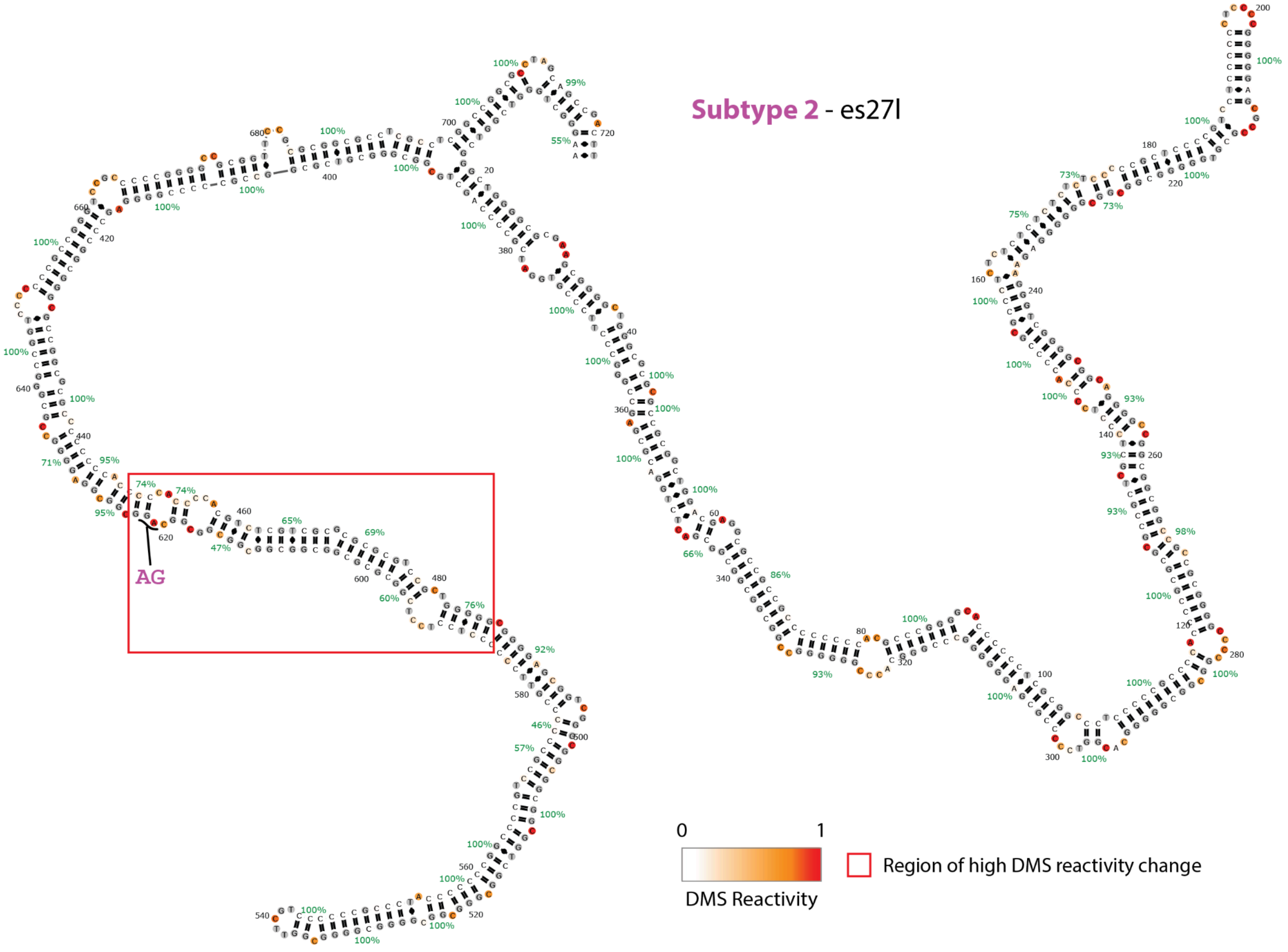


**Figure S20. In-cell DMS with long-read sequencing shows that es27l of different 28S subtypes have different RNA 2D structure**

RNA secondary structure of es15l predicted secondary structure for subtype 2 (G, AG, C).

A

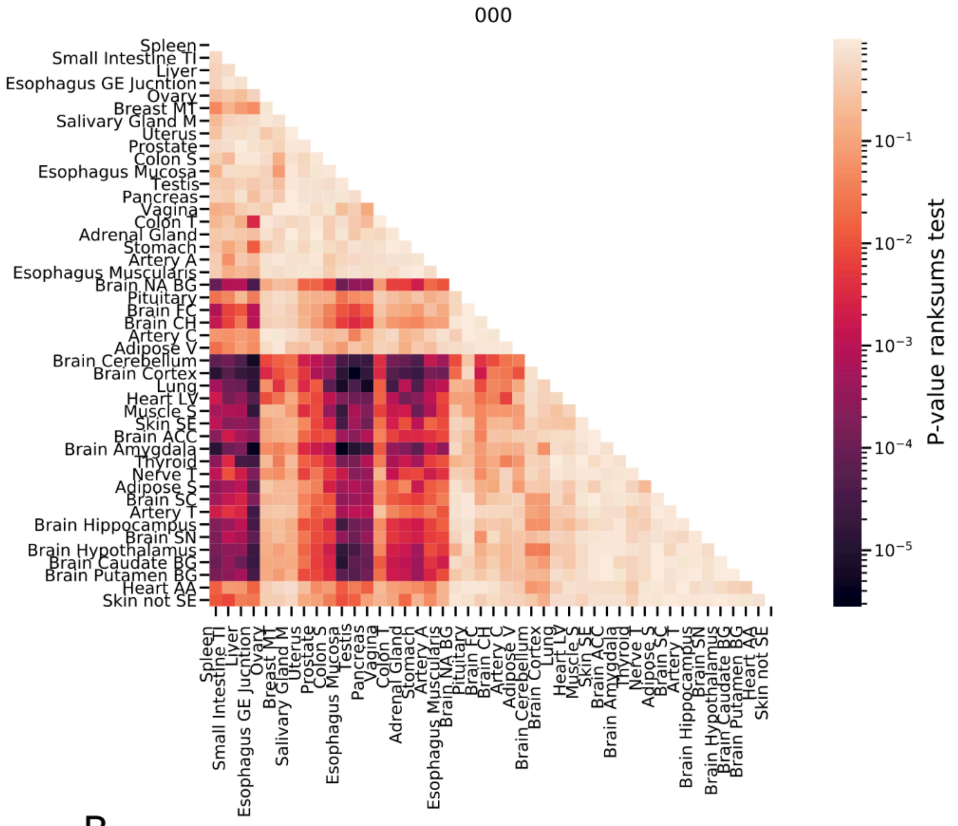
000
Spleen
Small Intestine TI
Liver
Esophagus GE Jucntion
Ovary
Breast MT
Salivary Gland M
Uterus
Prostate
Colon S
Esophagus Mucosa
Testis
Pancreas
Vagina
Colon T
Adrenal Gland
Stomach
Artery A
Esophagus Muscularis
Brain NA BG
Pituitary
Brain FC
Brain CH
Artery C
Adipose V
Brain Cerebellum
Brain Cortex
Lung
Heart LV
Muscle S
Skin SE
Brain ACC
Brain Amygdala
Thyroid
Nerve T
Adipose S
Brain SC
Artery T
Brain Hippocampus
Brain SN
Brain Hypothalamus
Brain Caudate BG
Brain Putamen BG
Heart AA
Skin not SE
$10^{-1}$
$10^{-2}$
$10^{-3}$
$10^{-4}$
$10^{-5}$
P-value ranksums test

B

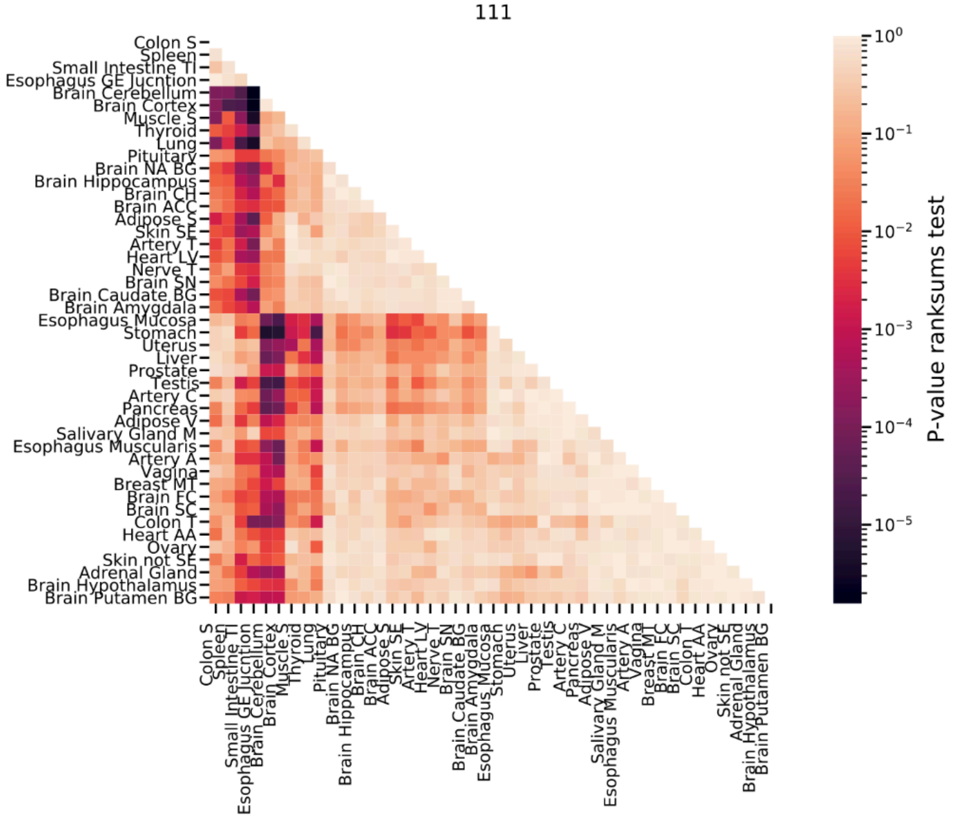
111
Colon S
Spleen
Small Intestine TI
Esophagus GE Jucntion
Brain Cerebellum
Brain Cortex
Muscle S
Thyroid
Lung
Pituitary
Brain NA BG
Brain Hippocampus
Brain CH
Brain ACC
Adipose S
Skin SE
Artery T
Heart LV
Nerve T
Brain SN
Brain Caudate BG
Brain Amygdala
Esophagus Mucosa
Stomach
Uterus
Liver
Prostate
Testis
Artery C
Pancreas
Adipose V
Salivary Gland M
Esophagus Muscularis
Artery A
Vagina
Breast MT
Brain FC
Brain SC
Colon T
Heart AA
Ovary
Skin not SE
Adrenal Gland
Brain Hypothalamus
Brain Putamen BG
$10^{0}$
$10^{-1}$
$10^{-2}$
$10^{-3}$
$10^{-4}$
$10^{-5}$
P-value ranksums test

**Figure S21 (1 out of 5 similar plots). rRNA subtype expression levels are tissue specific**

A. A heatmap showing FDR-corrected rank sum test P-values comparing the expression levels of the rRNA subtype with the haplotype sequence of G,AG,C (titled 000) across tissues**.** The tissues are ordered by average hierarchical clustering of the rank sum corrected P-values.
B. Same as (A) for haplotype A,GGCAG,T (titled 111) across tissues**.**

A

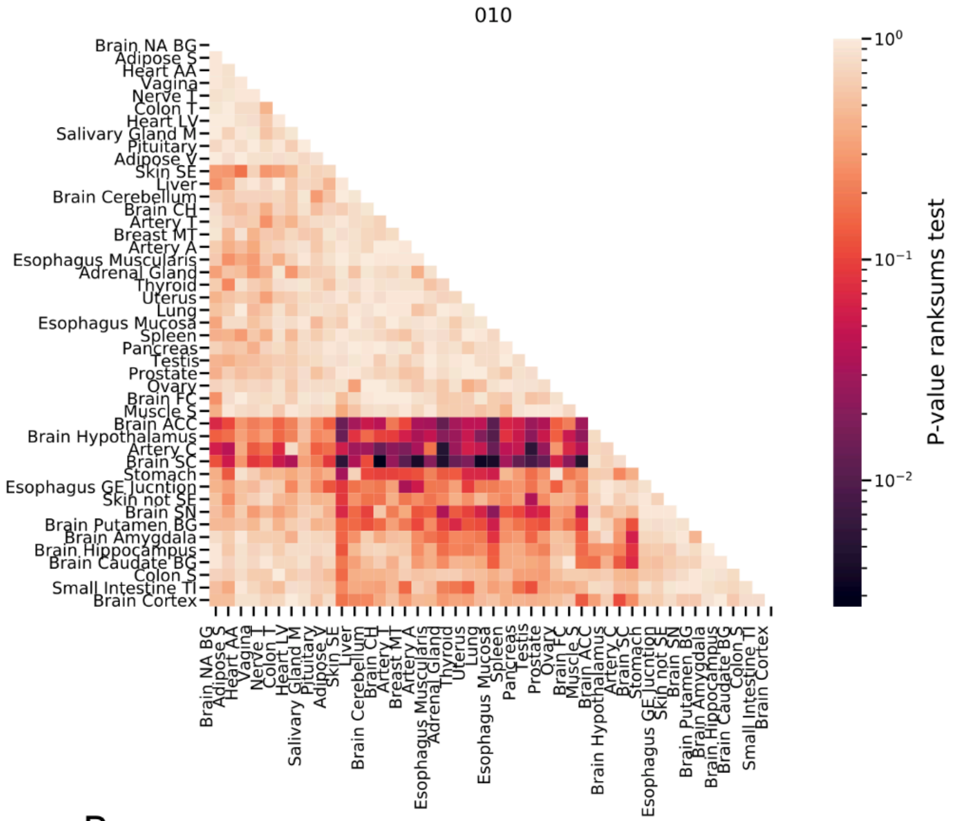
010
Brain NA BG
Adipose S
Heart AA
Vagina
Nerve T
Colon T
Heart LV
Salivary Gland M
Pituitary
Adipose V
Skin SE
Liver
Brain Cerebellum
Brain CH
Artery T
Breast MT
Artery A
Esophagus Muscularis
Adrenal Gland
Thyroid
Uterus
Lung
Esophagus Mucosa
Spleen
Pancreas
Testis
Prostate
Ovary
Brain FC
Muscle S
Brain ACC
Brain Hypothalamus
Artery C
Brain SC
Stomach
Esophagus GE Jucntion
Skin not SE
Brain SN
Brain Putamen BG
Brain Amygdala
Brain Hippocampus
Brain Caudate BG
Colon S
Small Intestine TI
Brain Cortex
$10^0$
$10^{-1}$
$10^{-2}$
P-value ranksums test

B

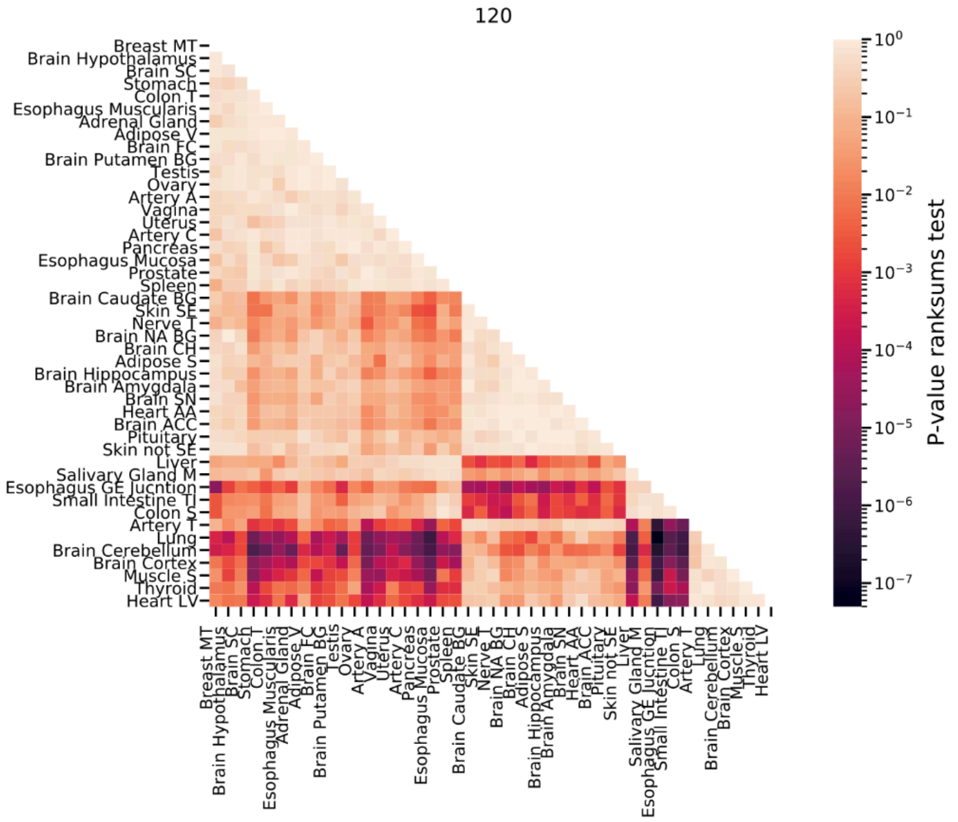
120
Breast MT
Brain Hypothalamus
Brain SC
Stomach
Colon T
Esophagus Muscularis
Adrenal Gland
Adipose V
Brain FC
Brain Putamen BG
Testis
Ovary
Artery A
Vagina
Uterus
Artery C
Pancreas
Esophagus Mucosa
Prostate
Spleen
Brain Caudate BG
Skin SE
Nerve T
Brain NA BG
Brain CH
Adipose S
Brain Hippocampus
Brain Amygdala
Brain SN
Heart AA
Brain ACC
Pituitary
Skin not SE
Liver
Salivary Gland M
Esophagus GE Jucntion
Small Intestine TI
Colon S
Artery T
Lung
Brain Cerebellum
Brain Cortex
Muscle S
Thyroid
Heart LV
$10^0$
$10^{-1}$
$10^{-2}$
$10^{-3}$
$10^{-4}$
$10^{-5}$
$10^{-6}$
$10^{-7}$
P-value ranksums test

**Figure S22 (2 out of 5 similar plots). rRNA subtype expression levels are tissue specific**

A. A heatmap showing FDR-corrected rank sum test P-values comparing the expression levels of the rRNA subtype with the haplotype sequence of G,GGCAG,C (titled 010) across tissues**.** The tissues are ordered by average hierarchical clustering of the rank sum corrected P-values.
B. Same as (A) for haplotype A,GG,C (titled 120) across tissues**.**

A

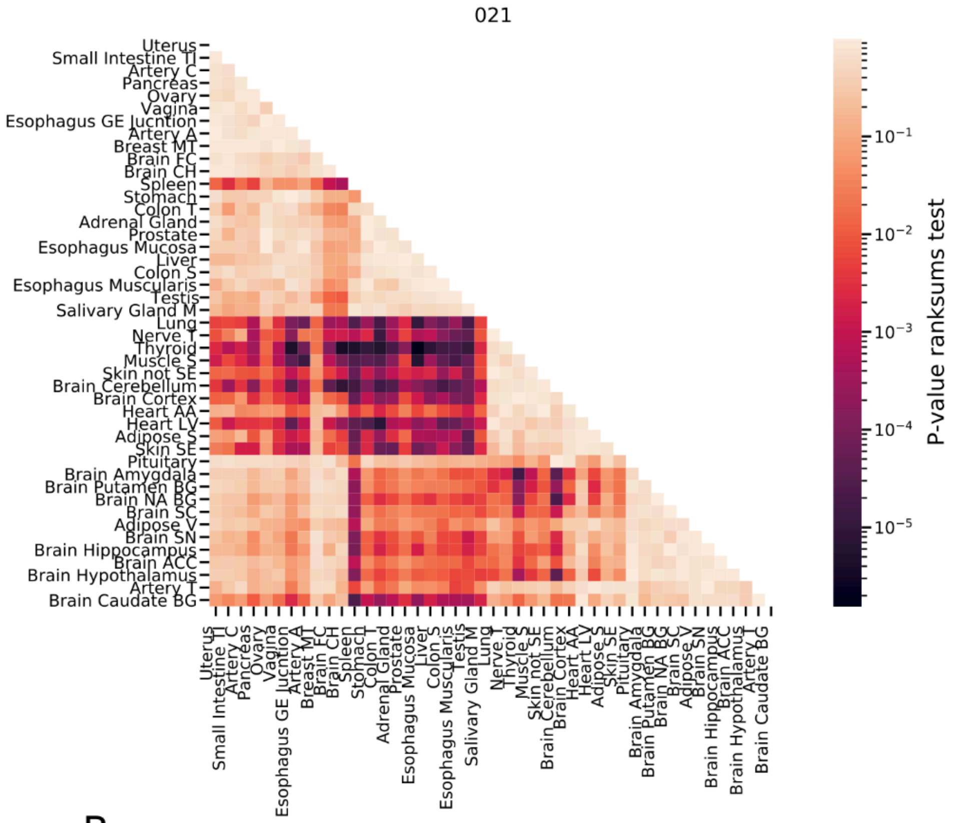
021
Uterus
Small Intestine TI
Artery C
Pancreas
Ovary
Vagina
Esophagus GE Junction
Artery A
Breast MT
Brain FC
Brain CH
Spleen
Stomach
Colon T
Adrenal Gland
Prostate
Esophagus Mucosa
Liver
Colon S
Esophagus Muscularis
Testis
Salivary Gland M
Lung
Nerve T
Thyroid
Muscle S
Skin not SE
Brain Cerebellum
Brain Cortex
Heart AA
Heart LV
Adipose S
Skin SE
Pituitary
Brain Amygdala
Brain Putamen BG
Brain NA BG
Brain SC
Adipose V
Brain SN
Brain Hippocampus
Brain ACC
Brain Hypothalamus
Artery T
Brain Caudate BG
$10^{-1}$
$10^{-2}$
$10^{-3}$
$10^{-4}$
$10^{-5}$
P-value ranksums test

B

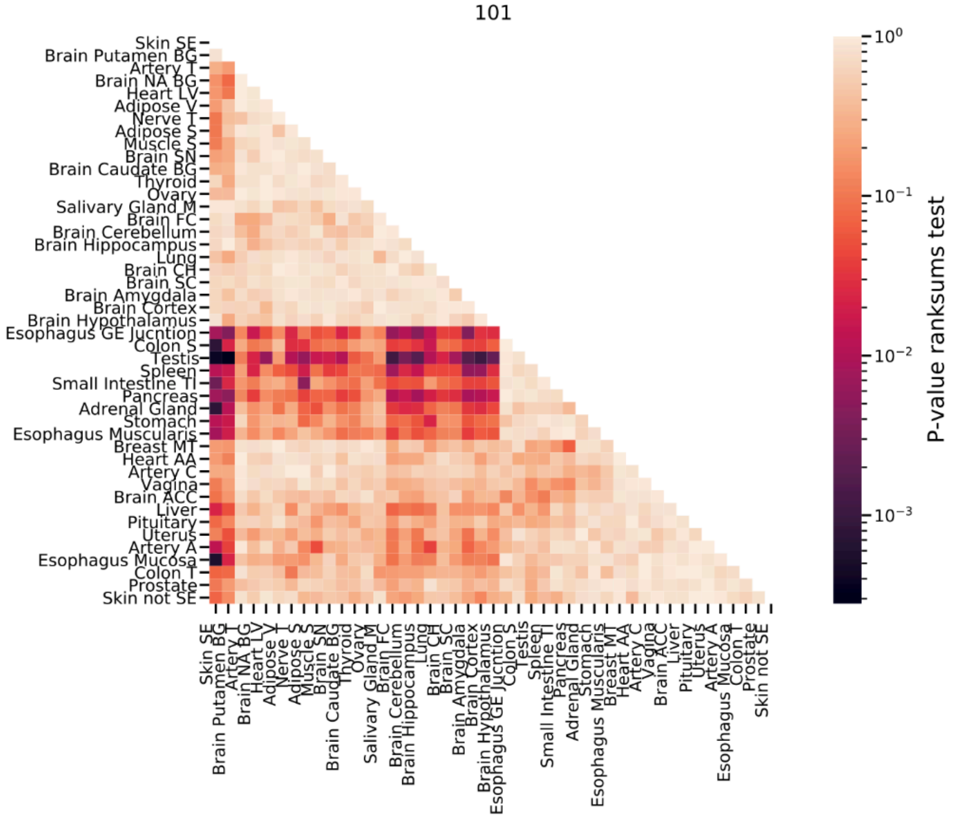
101
Skin SE
Brain Putamen BG
Artery T
Brain NA BG
Heart LV
Adipose V
Nerve T
Adipose S
Muscle S
Brain SN
Brain Caudate BG
Thyroid
Ovary
Salivary Gland M
Brain FC
Brain Cerebellum
Brain Hippocampus
Lung
Brain CH
Brain SC
Brain Amygdala
Brain Cortex
Brain Hypothalamus
Esophagus GE Junction
Colon S
Testis
Spleen
Small Intestine TI
Pancreas
Adrenal Gland
Stomach
Esophagus Muscularis
Breast MT
Heart AA
Artery C
Vagina
Brain ACC
Liver
Pituitary
Uterus
Artery A
Esophagus Mucosa
Colon T
Prostate
Skin not SE
$10^{0}$
$10^{-1}$
$10^{-2}$
$10^{-3}$
P-value ranksums test

**Figure S23 (3 out of 5 similar plots). rRNA subtype expression levels are tissue specific**

A. A heatmap showing FDR-corrected rank sum test P-values comparing the expression levels of the rRNA subtype with the haplotype sequence of G,GG,C (titled 021) across tissues**.** The tissues are ordered by average hierarchical clustering of the rank sum corrected P-values.
B. Same as (A) for haplotype A,AG,T (titled 101) across tissues**.**

A

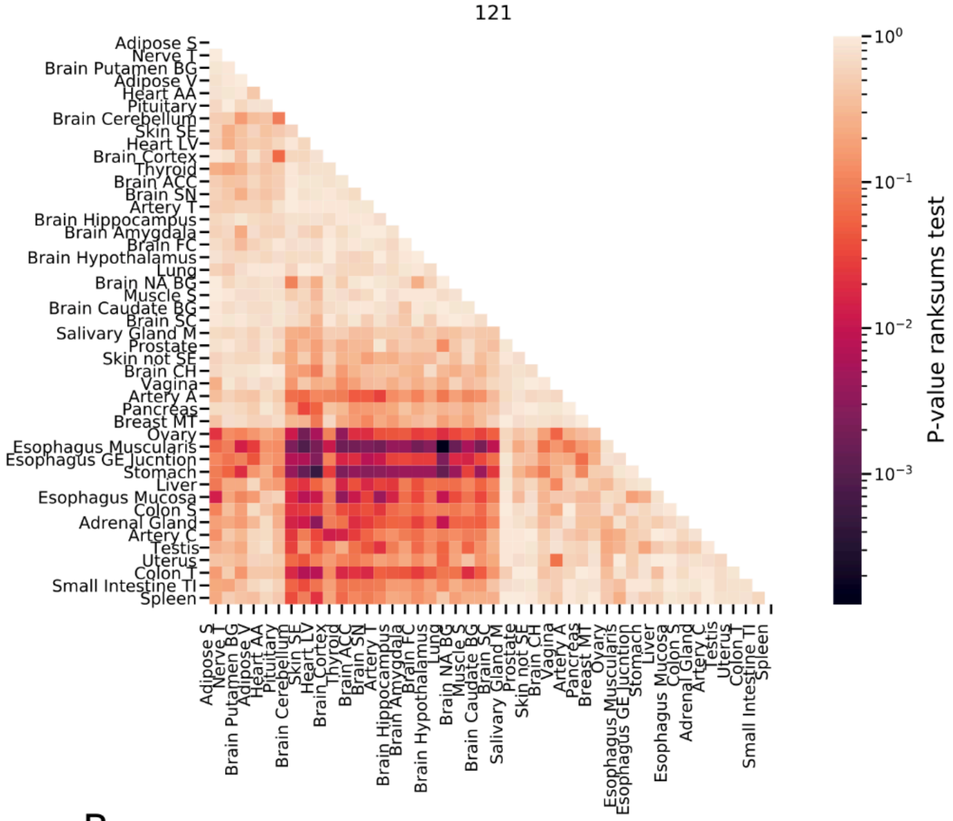
121
Adipose S
Nerve T
Brain Putamen BG
Adipose V
Heart AA
Pituitary
Brain Cerebellum
Skin SE
Heart LV
Brain Cortex
Thyroid
Brain ACC
Brain SN
Artery T
Brain Hippocampus
Brain Amygdala
Brain FC
Brain Hypothalamus
Lung
Brain NA BG
Muscle S
Brain Caudate BG
Brain SC
Salivary Gland M
Prostate
Skin not SE
Brain CH
Vagina
Artery A
Pancreas
Breast MT
Ovary
Esophagus Muscularis
Esophagus GE Jucntion
Stomach
Liver
Esophagus Mucosa
Colon S
Adrenal Gland
Artery C
Testis
Uterus
Colon T
Small Intestine TI
Spleen
$10^0$
$10^{-1}$
$10^{-2}$
$10^{-3}$
P-value ranksums test

B

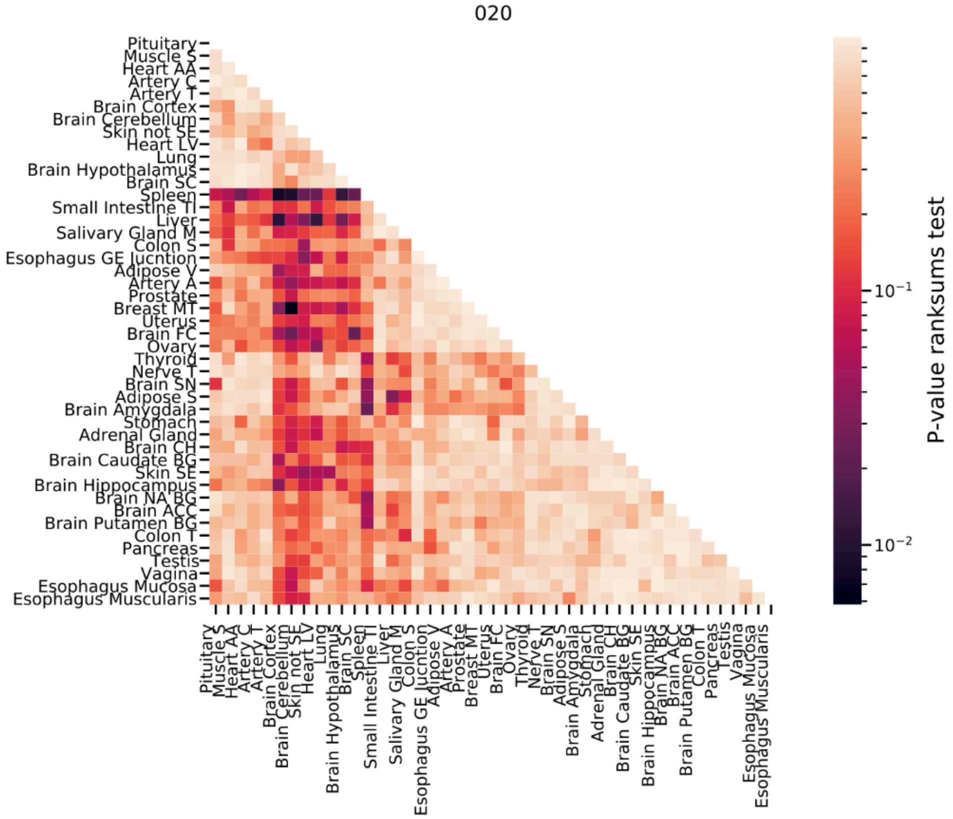
020
Pituitary
Muscle S
Heart AA
Artery C
Artery T
Brain Cortex
Brain Cerebellum
Skin not SE
Heart LV
Lung
Brain Hypothalamus
Brain SC
Spleen
Small Intestine TI
Liver
Salivary Gland M
Colon S
Esophagus GE Jucntion
Adipose V
Artery A
Prostate
Breast MT
Uterus
Brain FC
Ovary
Thyroid
Nerve T
Brain SN
Adipose S
Brain Amygdala
Stomach
Adrenal Gland
Brain CH
Brain Caudate BG
Skin SE
Brain Hippocampus
Brain NA BG
Brain ACC
Brain Putamen BG
Colon T
Pancreas
Testis
Vagina
Esophagus Mucosa
Esophagus Muscularis
$10^{-1}$
$10^{-2}$
P-value ranksums test

**Figure S24 (4 out of 5 similar plots). rRNA subtype expression levels are tissue specific**

A. A heatmap showing FDR-corrected rank sum test P-values comparing the expression levels of the rRNA subtype with the haplotype sequence of A,GG,T (titled 121) across tissues. The tissues are ordered by average hierarchical clustering of the rank sum corrected P-values.

B. Same as (A) for haplotype G,GG,C (titled 020) across tissues.

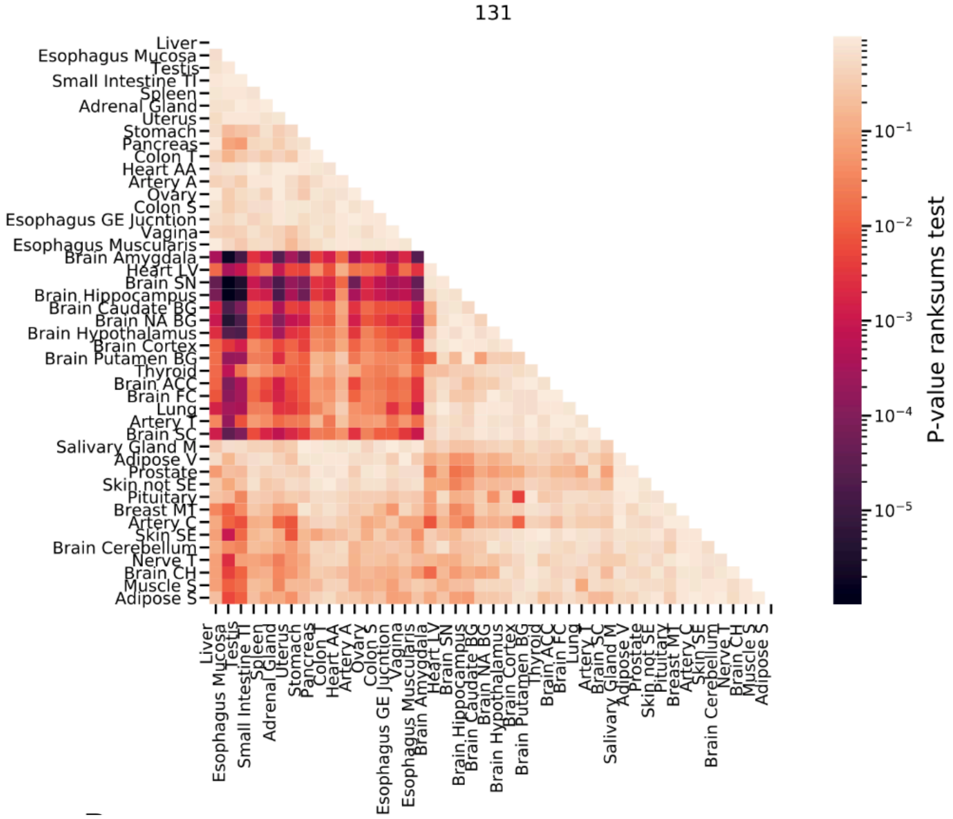


**Figure S25 (5 out of 5 similar plots). rRNA subtype expression levels are tissue specific**

A heatmap showing FDR-corrected rank sum test P-values comparing the expression levels of the rRNA subtype with the haplotype sequence of A,GGCGGCAG,T (titled 131) across tissues. The tissues are ordered by average hierarchical clustering of the rank sum corrected P-values.

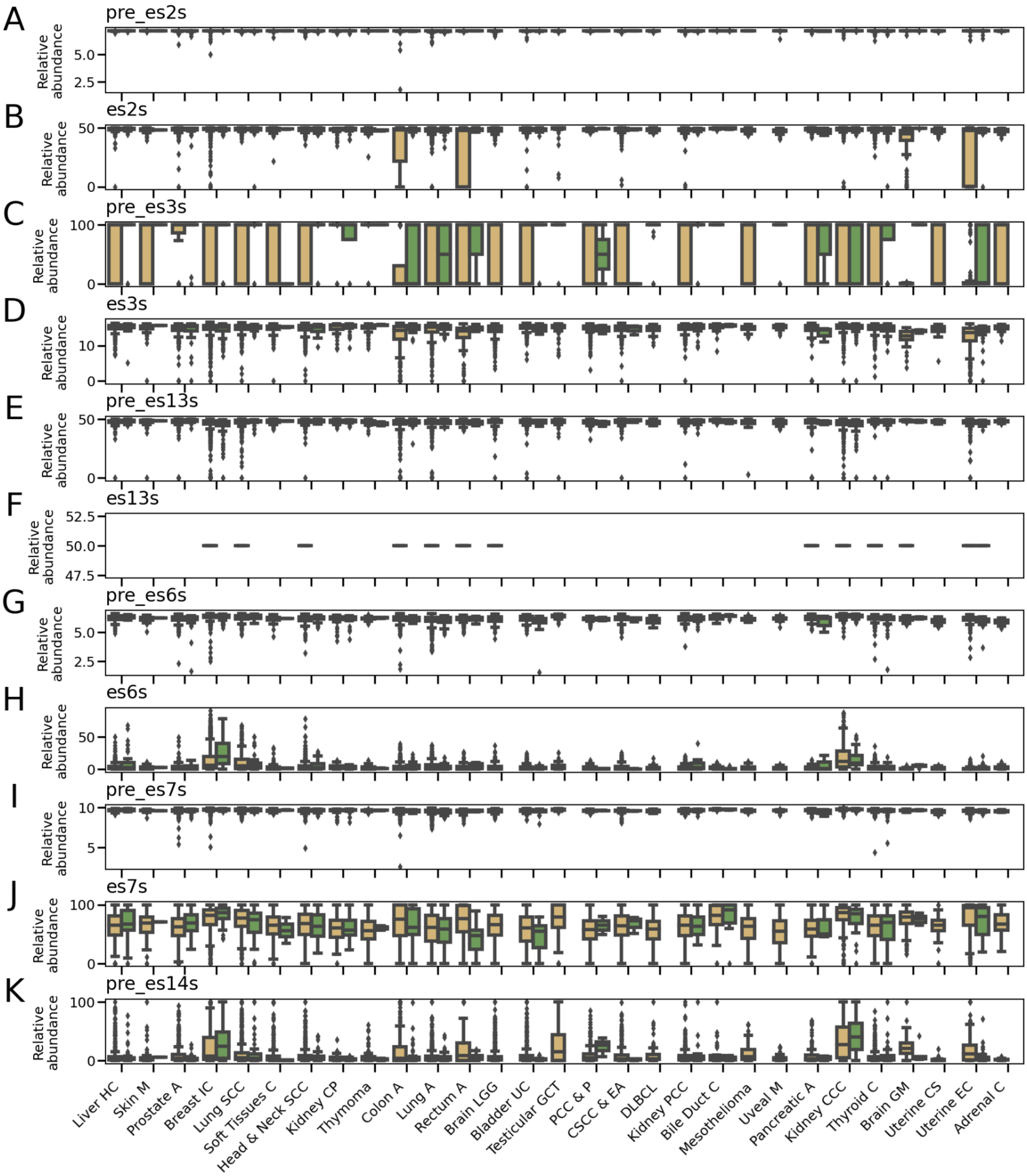


**Figure S26. Cancer-specific rRNA variants relative-abundances (1 of 6 figures)**

(A-K) Scatter plot of top abundant regional rRNA variants relative abundances for TCGA cancer and control biopsy samples (cancer and control samples are in yellow and green boxes respectively **Table**

**S23** for region to regional variant conversion and the P-value for comparing case/control). The top most abundant rRNA regional variant is presented per ES/non-ES region across tissues. The x-axis is the same for all panels and is displayed in (K).

Abbreviations:

Adrenal C = Adrenocortical Carcinoma
Bile Duct C = Cholangiocarcinoma
Bladder UC = Bladder Urothelial Carcinoma
Brain GM = Brain Glioblastoma Multiforme
Brain LGG = Brain Lower Grade Glioma
Breast IC = Breast Invasive Carcinoma
Colon A = Colon Adenocarcinoma
CSCC & EA = Cervical Squamous Cell Carcinoma and Endocervical Adenocarcinoma
DLBCL = Lymphoid Neoplasm Diffuse Large B-cell Lymphoma
Head & Neck SCC = Head and Neck Squamous Cell Carcinoma
Kidney CCC = Kidney Renal Clear Cell Carcinoma
Kidney CP = Kidney Chromophobe
Kidney PCC = Kidney Renal Papillary Cell Carcinoma
Liver HC = Liver Hepatocellular Carcinoma
Lung A = Lung Adenocarcinoma
Lung SCC = Lung Squamous Cell Carcinoma
Pancreatic A = Pancreatic Adenocarcinoma
PCC & P = Pheochromocytoma and Paraganglioma
Prostate A = Prostate Adenocarcinoma
Rectum A = Rectum Adenocarcinoma
Skin M = Skin Cutaneous Melanoma
Soft Tissues C = Soft Tissues Carcinoma
Testicular GCT = Testicular Germ Cell Tumors
Thyroid C = Thyroid Carcinoma
Uterine CS = Uterine Carcinosarcoma
Uterine EC = Uterine Corpus Endometrial Carcinoma
Uveal M = Uveal Melanoma

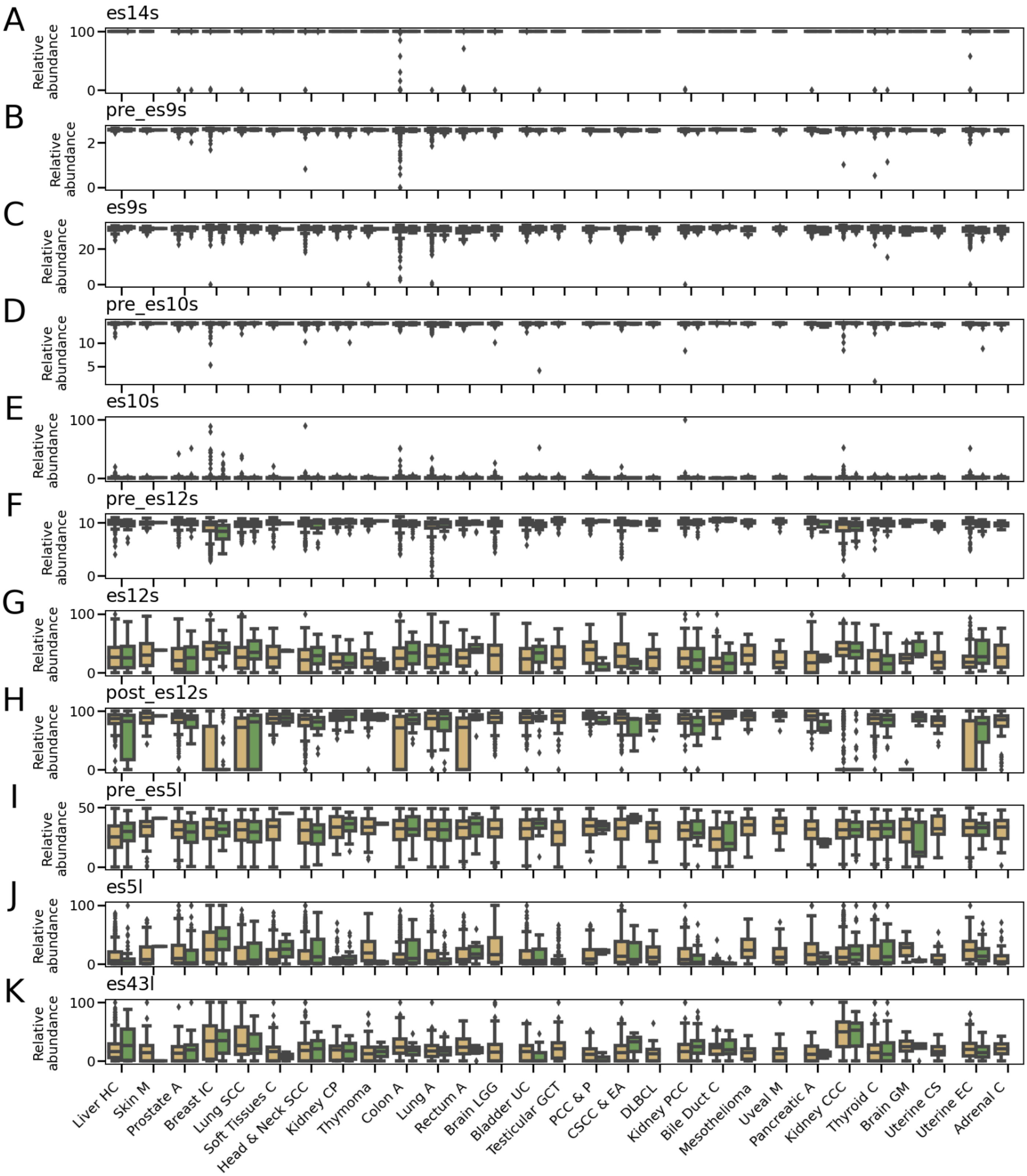


**Figure S27. Cancer-specific rRNA variants relative-abundances (2 of 6 figures)**

(A-K) Scatter plot of different regional rRNA variants, relative abundances for TCGA cancer biopsy samples (cancer and control samples are in yellow and green boxes respectively). The top most

abundant rRNA regional variant is presented per ES/non-ES region across tissues (**Table S22** for region to regional variant conversion and the P-value for comparing case/control). The x-axis is the same for all panels and is displayed in (K). X-axis cancer type full name for the abbreviations are listed at the bottom of **Figure S26**.

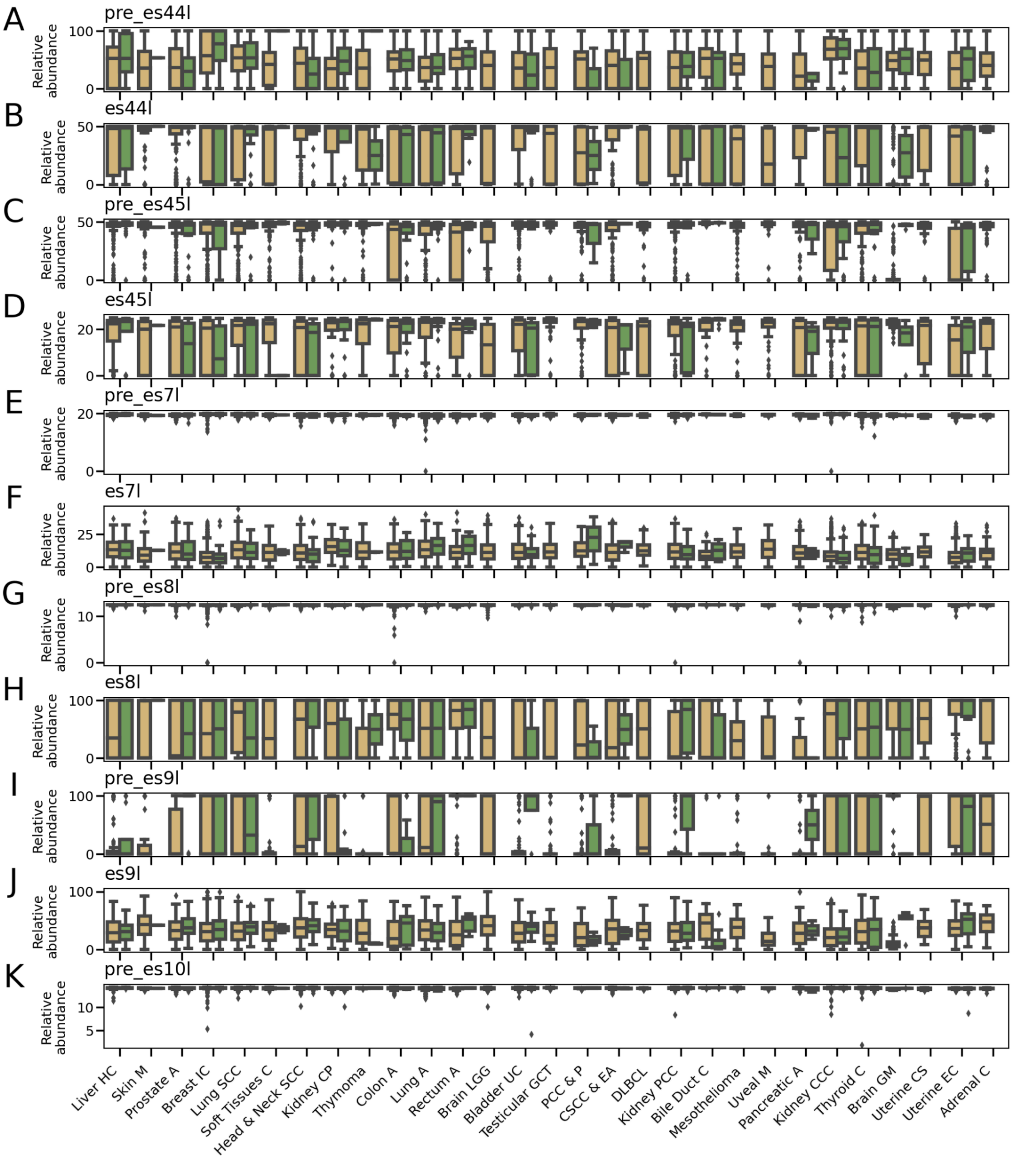


**Figure S28. Cancer-specific rRNA variants relative-abundances (3 of 6 figures)**

(A-K) Scatter plot of different regional rRNA variants, relative abundances for TCGA cancer biopsy samples (cancer and control samples are in yellow and green boxes respectively). The top most

abundant rRNA regional variant is presented per ES/non-ES region across tissues (**Table S23** for region to regional variant conversion and the P-value for comparing case/control). The x-axis is the same for all panels and is displayed in (K). X-axis cancer type full name for the abbreviations are listed at the bottom of **Figure S26**.

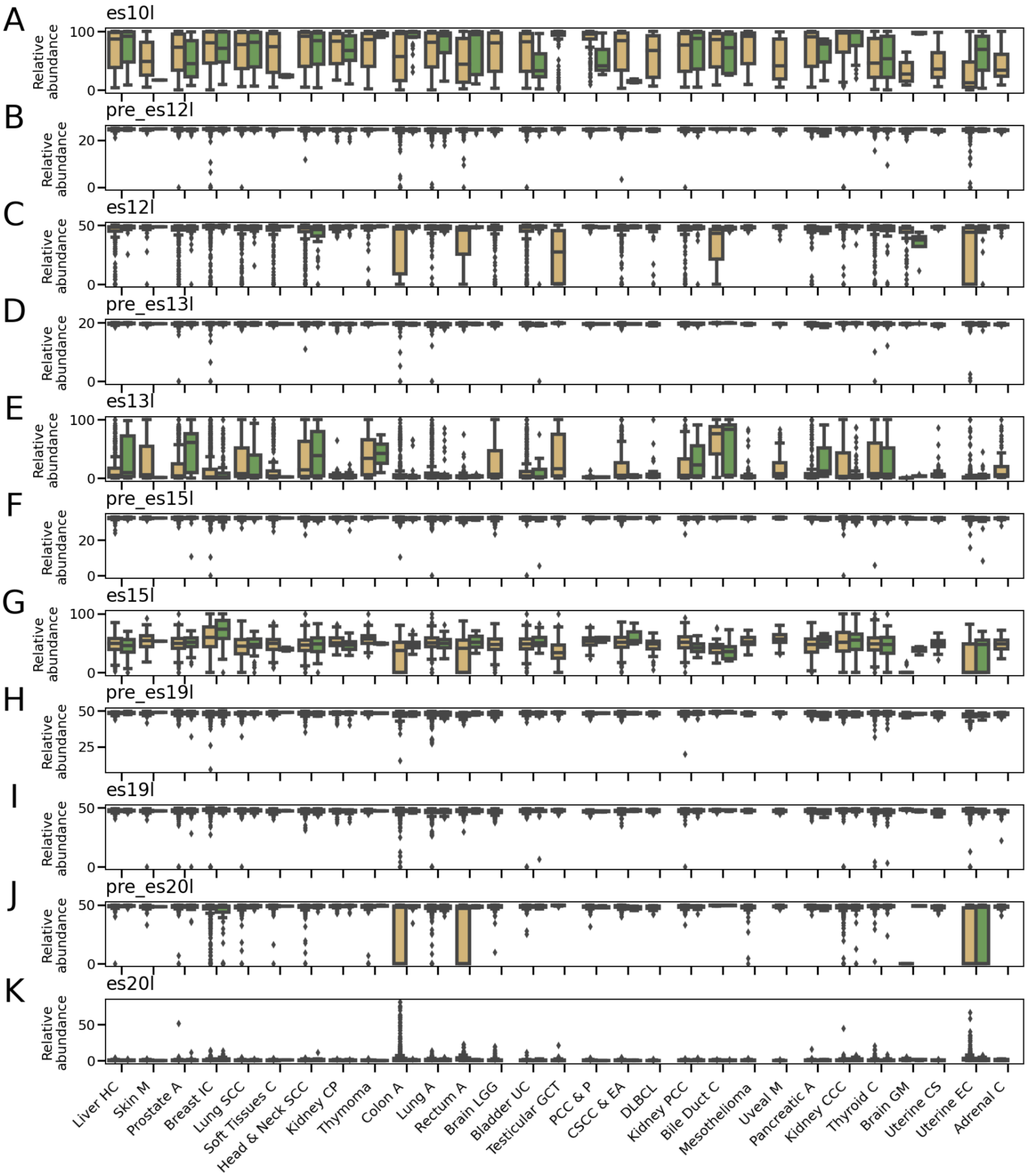


**Figure S29. Cancer-specific rRNA variants relative-abundances (4 of 6 figures)**

(A-K) Scatter plot of different regional rRNA variants, relative abundances for TCGA cancer biopsy samples (cancer and control samples are in yellow and green boxes respectively). The top most

abundant rRNA regional variant is presented per ES/non-ES region across tissues (**Table S23** for region to regional variant conversion and the P-value for comparing case/control). The x-axis is the same for all panels and is displayed in (K). X-axis cancer type full name for the abbreviations are listed at the bottom of **Figure S26**.

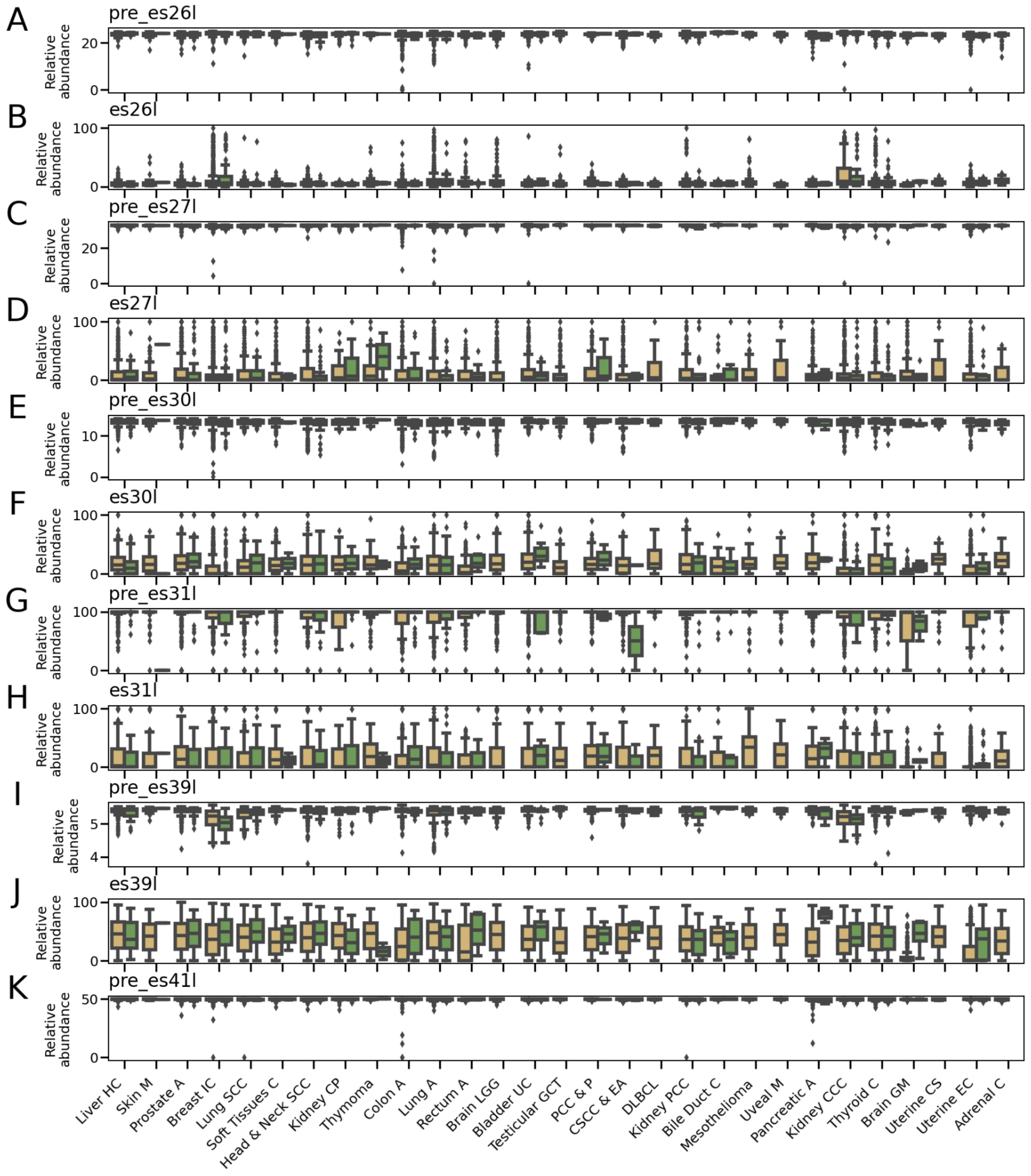


**Figure S30. Cancer-specific rRNA variants relative-abundances (5 of 6 figures)**

(A-K) Scatter plot of different regional rRNA variants, relative abundances for TCGA cancer biopsy samples (cancer and control samples are in yellow and green boxes respectively). The top most

abundant rRNA regional variant is presented per ES/non-ES region across tissues (**Table S23** for region to regional variant conversion and the P-value for comparing case/control). The x-axis is the same for all panels and is displayed in (K). X-axis cancer type full name for the abbreviations are listed at the bottom of **Figure S26**.

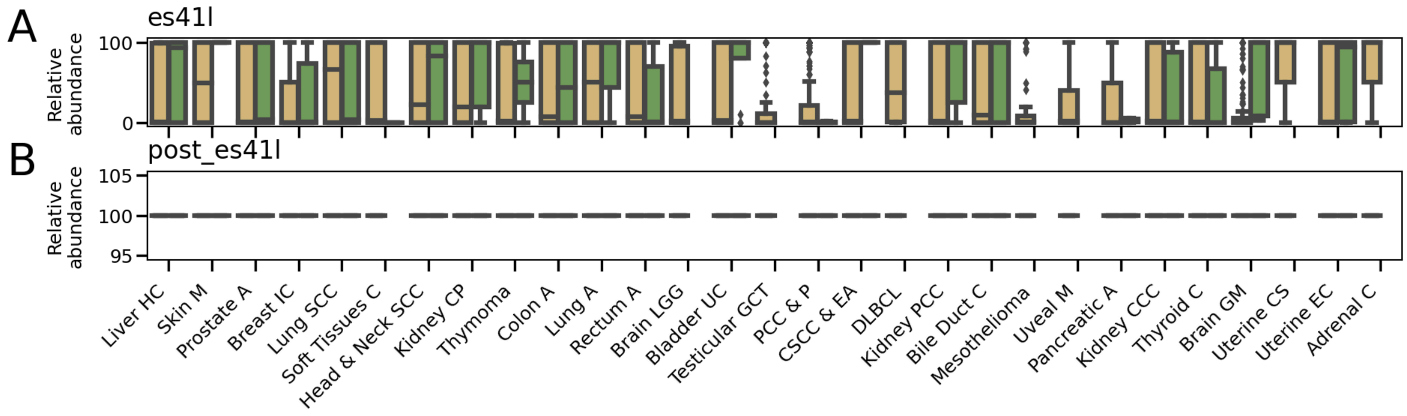


**Figure S31. Cancer-specific rRNA variants relative-abundances (6 of 6 figures)**

(A-B) Scatter plot of different regional rRNA variants, relative abundances for TCGA cancer biopsy samples (cancer and control samples are in yellow and green boxes respectively). The top most abundant rRNA regional variant is presented per ES/non-ES region across tissues (**Table S23** for region to regional variant conversion and the P-value for comparing case/control). The x-axis is the same for all panels and is displayed in (B).  X-axis cancer type full name for the abbreviations are listed at the bottom of **Figure S26**.

### Experimental Procedures

#### Reference index for rDNA extraction

For calling variants we map against three rDNA references :

1. Hg38 Un_GL000220v1 positions:105,423-118,723
2. A consensus from mapped Hg38 regions to Un_GL000220v1:105,423-118,723
3. A consensus rDNA from T2T v1.0 assembly of CHM13 created by multiple sequence alignment using Clustal Omega[1,2]

#### 18S and 28S extraction from hESC, GIAB and T2T, 1KGP

rDNA calling - hESC and GIAB: Reads were extracted by mapping HiFi fastq long reads (**Table S1, S24** for HiFi sample list) to the “Reference index for rDNA extraction” using minimap2[3] with “-N 20 -ax map-ont” parameters and processes with samtools[4].
T2T rDNA calling: We used complete 219 rDNA copies with 18S and 28S annotation by T2T.
hESC rRNA calling: The RT with 3’ primers of 18S and 28S results in 18S and 28S rRNA reads.

For both rDNA and rRNA, we keep reads that we consider full or near full length as followed:

1. We split long reads to consecutive non-overlapping 50bp short reads and map the short reads to “Reference index for rDNA extraction” using bowtie2 using default parameters[5].
2. To call a long read 18S, we demand a long read to have at least 18 short reads (which is the equivalent of ~900bp) to map to the 18S gene. For 28S calling, we demand a read to have at least 50 short reads (which is the equivalent of ~2500bp) to map to the 28S gene.
3. We keep 18S reads in the length range 1,500-2,100 bp and 28S reads in the length range 4,500-5,500 bp.
4. For calling deletion variants, in order to avoid calling variants where the RT stopped, we only considered reads that do not have deletions at the beginning at position 56 in the 18S and position 25 in the 28S and only report deletion atlas variants after these positions.

#### Reference Gap Alignment (RGA) method

Python implementation source code is available here:
https://github.com/daphnar/rRNA

1. We classified sequences as either 18S or 28S followed by Needleman–Wunsch global sequence alignment [6] of each sequence to one RNA45S5 reference (either 18S or 28S based on read classification) [7].
2. We created a reference sequence that aligns with all other sequences that we call a gap-aligned reference. This gap-aligned reference has the same sequence as the reference, but at each nucleotide position, we extended a gap at the size of the maximal gap found by the global sequence alignment to all sequences. Importantly, this gap-aligned reference allows straightforward comparison among all sequences without requiring computationally expensive all-by-all pairwise sequence alignments.
3. We aligned all H7-hESC sequences to the gap-aligned reference using the previous global alignment with additional extended gaps at reference positions.
4. Lastly, we extracted all variants at a given position across all aligned sequences.

Notably, we have benchmarked the gap-penalty opening and extension which can affect the indel number (**Table S4**). Since benchmarked parameters yielded a similar total number of indels, we use the default Needleman–Wunsch parameters of high penalty of gap opening and low penalty of gap extension.

#### Nucleotide variant calling in long reads

We ran our four step RGA, algorithm on the reads that pass the criteria in "18S and, 28S extraction from 1KGP,  GIAB, hESC, and T2T"

The output of this alignment are exact alignment of all reads against the 18S/28S reference. With this we extract all sequence variants of the 18S and 28S both in rRNA and rDNA.

For the 1KGP dataset, we report rDNA variants that were found in at least 5 reads and detected in 3 samples in **Table S5** (out of 30 1KGP samples with HiFi reads). For GIAB 2 trio families, we report variants found in at least 5 reads and detected in 2 samples **Table S6-S7** (For HiFi and ONT datasets). Since the Chinese Han father sample was not sequenced in ONT, we did not include this sample in the HiFi dataset and the total number of samples were 5: Ashkenazi mother, father, son, and Chinese Han mother and son.

#### Atlas variant calling in long-reads

We ran “Nucleotide, helix and ES variant calling” on both the hESC rDNA and the rRNA data and call a found variant an atlas variant if the variant is present in abundance greater than the HiFi read accuracy and are found in both rDNA and rRNA.
HiFi accuracy is expected to be greater than 99.9%.

For the hESC we obtained the following from “18S and, 28S extraction from hESC, GIAB and T2T“ step:
From the hESC rDNA we have obtained 762 complete 18S sequences and 386 complete 28S sequences.
From rRNAs, we have obtained 7,454 and 51,040 complete 18S sequences from monosome and polysomes, and 5,834 and 8,596 complete 28S sequences from monosome and polysomes.

Then, assuming HiFi accuracy of 99.9%, we call atlas variants that satisfy:

A. Nucleotide variant was found in the rDNA at least twice.
B. Nucleotide variant abundance in rRNA is at least 59 for the 18S and 10 for 28S.

The raw read count for nucleotide variants are reported in the atlas. After finding nucleotide atlas variants, to call for atlas helix and ES variants, we use the helix/ES annotation (**Table S13,14**) to aggregate nucleotide atlas variants at a given region and consider variants if they are found in both rDNA and rRNA. The raw read count for helix and ES resolution atlas is found in the names of the variants. There, the variant name ID indicates in the name the raw rDNA and rRNA read count.
The naming convention is “atlas_resolution:regional_variant:ID_Raw-rDNA-count_Raw-rRNA-count”.
So for example, at the atlas resolution of expansion segments, the first regional variant of region es2s is named: ES:es2s:0_d730_r43137. In this example, this nucleotide sequence containing sequence variations was observed 730 in rDNA and 43,137 in rRNA.

#### Validation of atlas nucleotide variants using 1KGP short-read data

We use the common Bowtie2 mapper tool and map the short-reads data from the 30 individuals from the 1KGP for which we have long-read data and map short-reads to our atlas of expanded resolution which allows mapping short-reads against it. This atlas contains complete expansion segments and non-expansion segments (ES/Non-ES marked in yellow and purple in **Figure S11**) which we also extend by 100 bases of the reference sequence to allow mapping to region ends. After mapping to this atlas, we only consider perfect matched reads. Afterwards, for finding which indels and SNVs are detected, we convert variants found at ES resolution back to nucleotide variants.

#### In-cell DMS probing for long-read sequencing

Approximately 2 x 10^7 of H7-hESC were used for in-cell DMS probing. Cells were washed with pre-warmed DPBS (Gibco, 14040133) prior to dissociation with Accutase (Gibco, A1110501) for ~5 mins at 37 °C. Then, cells were neutralized with mTeSR1 (StemCell Technologies, 85850) supplemented with 1µM thiazovivin (Tocris, 3845) and pelleted down by centrifuging at 200 x g at room temperature for 3 mins. Cells were then resuspended in 2,800 µL of pre-warmed mTeSR1+Tv. Then, 800 µL of pre-warmed 1 M bicine (Fisher Scientific, ICN10100580) buffer (pH 8.3 at 37 °C) was added, followed by 400 µL of 16% DMS (Sigma Aldrich, D186309) diluted in 100% ethanol (Gold Shield Distributors, 0412804-PINT). DMS labelling was done by incubating the mixture at 37 °C for 5 mins, prior to be quenched by adding 2,000 µL of ice-cold BME (Sigma Aldrich, M3148). Cells were pelleted down by centrifuging at 200 x g at 4 °C for 3 mins, and then lysed by resuspending them in 8 mL of cold TRIzolTM reagent (Invitrogen, 15596026). Solution was left at room temperature for 5 mins prior to adding 600 µL chloroform (Fisher Scientific, C298-500). The tube was then shaken vigorously for 15 seconds or so and left at room temperature for 3 mins. The sample was then centrifuged at 21,000 x g for 15 mins at 4 °C. A total of 4,440 µL aqueous phase was extracted and 4,440 µL of 100% ethanol was added before subjecting them into further cleanup and DNAse digest using Zymo RNA Clean and Concentrator Kit-5 as elaborated in “Polysome RNA extraction”.

50 µg of total RNA was used across ten 100 µL reactions. RT was done as described in “rRNA reverse transcription” section, with a few modifications. After RT, 0.4x beads by volume were used to size select cDNA. Ten reactions were then pooled together, and its cDNA concentration measured. For the second-strand synthesis, each reaction was done with a maximum of 500 ng of cDNA. Afterwards, PacBio IsoSeq library was constructed as per described in “PacBio SMRT sequencing library preparation” section.

#### rRNA subtype DMS reactivity and structure calling

28S sequenced reads from “In-cell DMS probing for long-read sequencing” were binned intro rRNA subtypes as followed:

1. We ran RGA method on the DMS reads

2. We bin DMS reads to rRNA subtype groups based on the hESC subtypes nucleotide positions at 60, 2188, 3513, 4913
3. Per DMS read, at each nucleotide we mark sequence variants that are not in the atlas as modified.
4. For every rRNA subtype group, we calculate the  rRNA subtype group reactivity as the average modification per nucleotide position across binned DMS reads.
5. Next we use  90% winsorization to set the DMS reactivity values from 0 to 1.
6. Lastly, we use the Biers Matlab package with RNAstructure and Varna for plotting [8]-[9].

#### 28S full length sequence comparisons and visualization in PCoA

In the T2T genome we discovered that although there are only 62 reported rDNA variants, the high frequency rDNA variants in H7-hESC also appeared in high frequency in the T2T rDNA (**Figure S9A**, R=0.8 Pearson correlation). In the GIAB samples, like in the H7-hESC, we found hundreds of variants with high agreement between their frequencies and H7-hESC frequencies (**Figure S9B-D**). We analyzed the linkage of the same positions in the T2T and GIAB, and found as found in the H7-hESC that es7l positions have low linkage, and es15l, es27l and es39l have relatively higher linkage within each region (**Figure S10**).

We compare the pairwise-distances between all hESC/GIAB 28S separately using 6-mer word base comparison with Alfree tools [10]. Pairwise distances are then visualized by plotting the first two PCos of the and the Bray-Curtis PCoA (**Figure 3D**). Each 28S is colored by the haplotype of that 28S as defined by the 60, 3513 and 4913 positions.

#### Atlas relative abundance calling for RNA short-read datasets

Short read RNA-seq are mRNA targeted however we found that about 2% of reads mapped to rRNA in the GTEx and TCGA. For comparing relative abundance across samples, we rarefaction samples of GTEx dataset to 500,000 rRNA mapped reads and TCGA to 250,000 rRNA mapped reads and throw samples with less than 100,000 rRNA mapped reads.

For short-reads, we use Kallisto[11] tool for region relative abundance estimation in the following way:
Once made for all GTEx/TCGA samples. We create one Kallisto index[11] combining the 18S and 28S variants in our ES/non-ES atlas with expanded reference (**Extended data 5-6** with “expand_100bp” in the name of the file) using Kallisto default parameters.

Linux command line:

```
> cat Extended_Data5.atlas.ES.18S.expand_100bp.fa > extended_atlas.fa
> cat Extended_Data6.atlas.ES.28S.expand_100bp.fa >> extended_atlas.fa
> kallisto index -i atlas_rRNA extended_atlas.fa
```

This expanded reference version of the atlas is the same ES/non-ES atlas with additional flanking 100bp on both ends (3' and 5') of the relevant region with the reference sequence. With these expansions, short reads that map to the 5' or 3' end of a region are mapped to the variants (as opposed to unmapped without expansions).

Here we chose our ES/non-ES atlas reference, as ES/non-ES regions are longer than helices (**Table S14**) but using the helix expanded reference atlas would give the same results (**Extended data 3-4** with "expand_100bp" in the name of the file).

We quantify all-region abundances of a sample (in the example below named FASTQ-FILE) using Kallisto[11] with default parameters.

Linux command line:

```
> kallisto quant -i atlas_rRNA FASTQ-FILE -o OUTPUT_DIRECTORY
```

Then, to compare expression of a given ES/non-ES regional variant, we normalize read count by the region length and normalize to one every ES/non-ES region independently.

Python code:

```
abundance =
pd.read_csv(os.path.join(OUTPUT_DIRECTORY,'abundance.tsv'),sep='\t',index_
col=0)[['eff_length','tpm']]
abundance = abundance['tpm']/abundance['eff_length']
ra = []
for group, group_df in abundance.groupby(lambda x: x.split(':')[1]):
    ra.append(group_df/group_df.sum())
normalized_abundnces = pd.concat(ra)
```

normalized_abundnces in the above python code contains the relative abundances of variants after normalization by ES/non-ES region.

#### GTEx and TCGA sample handling:

GTEx: Most individuals have multiple organs sequenced. To control for inter-individual variations when comparing tissues, we select one sample per individual in the GTEx dataset. For each compared tissue pair, individuals that have both tissues are randomly divided into two halves, from the first group we keep one tissue and from the second group we keep the other tissue. This way we only have one tissue per individual when comparing tissues. In all analyses we compared tissues with at least 10 samples.

In the TCGA cancer/control comparison, we selected cancers with at least 50 samples.

#### Polysome-RNA-extraction

H7-hESCs were harvested with Accutase (Gibco), and the cell pellets were lysed in lysis buffer (20 mM Tris pH 7.5, 150 mM NaCl, 15 mM $MgCl_2$, 100 µg/ml cycloheximide, 1 mM DTT, 0.5% Triton X-100, 0.1 mg/ml heparin, 8% glycerol, 20 U/ml TURBO DNase (Ambion, AM2238), 200 U/mL SUPERase In RNase Inhibitor (Ambion, AM2696), 1x Combined Protease and Phosphatase Inhibitor (Thermo Scientific, 78443)) at 4 ℃ for 30 minutes with occasional vortexing. Lysates were sequentially centrifuged at 1800g for 5 minutes at 4 ℃ and then at 10,000g for 5 minutes at 4 ℃, retaining the final supernatant as the cytoplasmic extract. Cytoplasmic extract was loaded on to a 10-45% sucrose gradient (20 mM Tris pH 7.5, 100 mM NaCl, 15 mM MgCl2, 100 µg/ml cycloheximide, made on a Biocomp Model 108 Gradient Master) and centrifuged in a Beckman SW41 rotor at 40,000 rpm for 2.5 hours at 4 ℃. Gradients were then fractionated on a Density Gradient Fraction System (Brandel, BR-188) with continuous A260 measurements. To each fraction (which contained approximately 750 µL), 100 µL of 10% SDS was added and the tubes vortexed to mix, followed by the addition of 140 µL of 3 M sodium acetate (pH 5.5) and 200 µL of RNase-free water, vortexing to mix. For ribosomal populations that spanned multiple fractions, such as the polysomes, equal volumes of each corresponding fraction was pooled in a separate tube to a total volume of 900 µL. To 900 µL of fractionated sample, 900 µL of acid phenol chloroform was added and heated at 65 ℃ for 5 minutes. The samples were then centrifuged at 21000g for 10 minutes at room temperature, and the aqueous phase transferred to a new tube. The aqueous phase was mixed with an equal volume of 100% ethanol and the RNA purified using the Zymo RNA Clean and Concentrator Kit-5 following manufacturer's instructions. DNase treatment was performed using TURBO DNase (1 µL of 2 U/µL per 50 µL reaction) at 37 ℃ for 30 minutes, and the RNA purified using the Zymo RNA Clean and Concentrator Kit-5 following manufacturer's instructions.

400 µL of cold cytoplasmic lysis buffer was added to each cell pellet. Cells were mixed and lysed by repeated vortexing for 30 s, followed by cooling down on ice for 30 s, repeated for 3 times in total. Cells were then incubated on ice for 30 minutes, vortexing for approximately 10 seconds every 10

minutes for complete lysis. Afterwards, cellular debris, organelles, and microsomes were removed with four serial centrifugations at 800 x g twice, 8,000 x g, and 21,300 x g for 5 mins each at 4 °C. RNA amount of the clarified cytoplasmic lysate was measured using nanodrop. Approximately 0.8-1 mg of RNA was set aside for sucrose gradient fractionation.

As the cells were being lysed, 10-45% sucrose gradient were prepared as follows: 50 mL of 10% and 45% sucrose buffers (20 mM Tris pH 7.5, 15 mM $MgCl_2$, 150 mM NaCl, 1 mM DTT, 100 µg/mL cycloheximide, 5 gr (10% solution) and 22.5 gr of sucrose (45% solution) (Millipore 8510-OP), in nuclease free water) were prepared separately. Using SW 41 Ti rotor compatible ultracentrifuge tubes (Beckman Coulter 331372), the two sucrose gradient buffers were layered extremely gently, and then the gradient was established using a gradient maker (Biocomp Gradient Master 108).

The cytoplasmic lysate was then layered on top of the sucrose gradient. The tubes were then loaded into SW 41 Ti rotor and centrifuged at 40,000 rpm for 2.5 hours at 4 °C. Afterwards, the gradient was then fractionated into 16 2 mL tubes every 30 seconds of ~700 µL solution each using a fractionation system (Brandel BR-188). The A260 trace was used as a reference to determine where the free ribonucleoproteins, free subunits, monosomes, and polysomes were. 100 µL of 10% SDS (Invitrogen AM9820) and 200 µL of 1.5 M sodium acetate (Invitrogen AM9740) were added into each fraction.

RNA was extracted from each fraction by adding 500 µL of acid-phenol:chloroform, pH 4.5 with IAA (125:24:1) (Invitrogen AM9722). The fractions were then incubated at 65 °C, 500 rpm thermomixer for 5 minutes. The RNA-containing aqueous phase, ~700 µL, was separated from the organic phase by centrifugation at 21,300 x g for 15 minutes at 4 °C. Further cleanup and trace DNA removal were done as described in the section "Whole-cell RNA extraction".

#### Polysome fractions collection

We collected and sequenced RNA from ribosome containing fractions: ribosomes (monosomes) and polysomes (**Figure S1** for H7-hESC A260 trace).

#### rRNA reverse transcription

Reverse transcription was done using TGIRT-III enzyme (InGex TGIRT50) with modified buffer and reaction conditions to increase enzyme processivity against highly structured and modified rRNA. To start, 4 µL of 100 µM pooled barcoded RT primers were added into 4.3 µL of 1 µg of RNA.

RNA-primer mix was then denatured at 65 °C for 5 minutes. 2.5 µL of 8x RT buffer (600 mM KCl (Invitrogen AM9640G), 160 mM Tris pH 7.5, 80 mM $MgCl_2$) was then added at 65 °C, and the reaction then cooled to 25 °C. Subsequently, 8.7 µL of enzyme mix (12.2% of PEG 5000, 12.2 mM of DTT, 2.44 M of Betaine, 4.88 U/µL of TGIRT-III, 12.2 U/µL of SUPERaseIn RNAse Inhibitor) was added into each reaction, followed by 30 minutes incubation at 25 °C. Afterwards, 1 µL of 25 mM dNTP was added prior to incubating the samples at 60 °C for 2 hours. The final concentration of the reagents in the 20 µL RT reaction are: 20 mM Tris HCl, 75 mM KCl, 10 mM $MgCl_2$, 5% PEG 5000, 5 mM DTT, 1 M Betaine, 2 U/µL TGIRT-III, 1 U/µL SUPERaseIN RNAse Inhibitor, and 10 µM pooled RT primers for 1 µg of RNA. After the RT, the RNA template is hydrolyzed by adding 1 µL of 2.5 M NaOH at 95 °C for 3 minutes. After cooling down to 4 °C, the reaction was neutralized by adding 1 µL of 2.5 M HCl and 1 µL of 500 mM Tris pH 7.5.

SPRISelect magnetic beads (Beckman Coulter B23319) were used for cDNA cleanup following the manufacturer's protocol. Beads were washed to remove contaminants that elute simultaneously with the DNA and interfere with polymerase binding in PacBio Sequel IIe system. In brief, for every 500 µL of SPRISelect beads in a low binding tube (Eppendorf 0030108442), the beads were centrifuged down at 21,300 x g for 1 minute. The tube was then placed in a magnetic rack, and the supernatant was aspirated and kept aside. The beads were then washed with 1 mL of nuclease-free water, vortexed, centrifuged at 21,300 x g for 1 min, and placed in a magnetic rack to remove the water wash. The wash was repeated a total of five times. Afterwards, the beads were washed similarly with 1 mL of Qiagen buffer EB (Qiagen 19086) instead. The beads were then resuspended with the original supernatant reserved previously, and could be kept at 4 °C for at most a week.

To clean up the single-stranded cDNA with the washed beads, a 2.2x beads-to-sample volume ratio was added. The beads then were washed with freshly prepared 85% ethanol for 1 minute while the tubes were attached on the magnetic rack. cDNA was then eluted in 40 µL of nuclease-free water for second-strand synthesis.

For each reaction, 1 µL of 12.5 mM dNTP (Thermo Scientific R0181) and 1 µL of 2 µM random hexamer (Thermo Scientific SO181) were added. Making sure that all components were kept on ice, 26 µL of nuclease-free water, and 4 µL of second-strand synthesis enzyme with 8 µL of 10x buffer (NEB E6111S) were added for a total of 80 µL reaction volume. The sample was then incubated at 16 °C in the thermocycler with the lid heating turned off. Double-stranded cDNA was then cleaned up using SPRIselect following the manufacturer's protocol, with a 0.82x beads-to-reaction ratio instead.

Washing was done twice with freshly prepared 85% ethanol, and the cDNA was then eluted in 50 µL of nuclease-free water. Cleanup was repeated once again with the same bead ratio to enrich for full-length reverse-transcribed rRNA molecules, with the final elution done in 30 µL of nuclease free water. cDNA concentration was then measured with qubit

#### PacBio SMRT sequencing library preparation

Multiplexed library was made as described in “Iso-Seq™ Express Template Preparation for Sequel® and Sequel II Systems”. cDNA amplification was skipped to prevent possible amplification bias against highly structured or repetitive sequences. In brief: equal amounts of cDNA from each barcode was pooled for a total of ~200 ng prior to DNA damage repair step. After pooling, cDNA was concentrated with 1x volume of SPRIselect beads and eluted in 48 µL of nuclease-free water. DNA damage repair, end-repair / A-tailing, overhang adapter ligation, and the final library cleanup were performed according to the protocol mentioned above, substituting the ProNex beads with the washed SPRIselect beads.

#### HeLa cell culture

The Human HeLa cell line was sourced from ATCC(CCL-2), and subsequent culturing was performed in DMEM supplemented with 10% FBS at 37°C and 5% $CO_2$.

#### In situ rRNA sequencing experimental procedure

Glass-bottom 12-well plates (Mattek, P12G-1.5-14-F) were treated as follows: Oxygen plasma treatment was applied for 5 mins (Anatech Barrel Plasma System, 100W, 40% $O_2$), followed by sequential incubation with 1% methacryloxypropyltrimethoxysilane (Bind-Silane, GE Healthcare 17-1330-01) 88% ethanol (VWR, 89125-170), 10% acetic acid (Sigma-Aldrich, A6283-100ML), and 1% $H_2O$ (Thermo Fisher Scientific, 10977023) at room temperature for 1 hour and 0.1 mg/mL Poly-D-lysine (Sigma-Aldrich, P7280-5X5MG) solution at room temperature for an additional hour. Micro cover glasses (Electron Microscope Sciences, 72226-01) underwent a pretreatment step with Gel Slick (Lonza, 50640) at room temperature for 15 mins and were then air-dried.

HeLa cells were cultured in treated 12-well plates, and after rinsing with 1× PBS (Thermo Fisher Scientific, 10010049), they were fixed with 1 mL of 1.6% PFA (Electron Microscope Sciences, 15710-S) in PBS buffer at room temperature for 15 mins. Following fixation, the cells underwent permeabilization by treatment with 1 mL of pre-chilled (-20°C) methanol (Sigma-Aldrich, 34860-1L-R) and incubation at -20°C for an hour. Thereafter, HeLa cells were transferred from the -20°C fridge to room temperature for 5 mins, and then washed twice with PBSTR (0.1% Tween-20 (Calbiochem, 655206), 0.1 U/µL RNaseOUT (Thermo Fisher Scientific, 10777019) in PBS) for 5 mins each.

For the reverse transcription (RT) process, primers were prepared by dissolving them at a concentration of 250 µM in ultrapure RNase-free water (Thermo Fisher Scientific, 10977023), followed by pooling. All probes were manufactured by Integrated DNA Technologies (IDT). The probe mixture was subjected to heating at 90°C for 5 mins, followed by cooling to room temperature. The samples were then treated with 300 µL of template switching mixture, which included 1× template switching buffer (New England Biolabs, M0466L), 250 µM dNTP (Invitrogen 100004893), 40 µM 5-(3-aminoallyl)-dUTP (Invitrogen AM8439), 2.5 µM RT primer, 0.4 U/µL RNaseOUT, 3.3 µM template switching oligo, and 1× template switching RT enzyme mix. This mixture was incubated at 4°C for 15 mins, followed by an overnight placement in a 42°C humidified oven with gentle shaking.

The following day, the samples underwent three washes with 500 µL PBST (0.1% Tween-20 in PBS) for 5 mins each. To cross-link cDNA molecules containing aminoallyl-dUTP, the specimens were incubated with 5 mM $BS(PEG)_9$ (Thermo Fisher Scientific, 21582) in PBST for 1 hour at room temperature, followed by a wash with PBST at room temperature for 5 mins. The cross-linking reaction was quenched by treating the samples with 0.1 M Glycine (Sigma-Aldrich, 50046-250G) in PBST at room temperature for 30 mins. To degrade residual RNA and generate single-stranded cDNA, the specimens were incubated for 2 hours at 37°C with an RNA digestion mixture, composed of 0.25 U/µL RNase H (New England Biolabs, M0297L), 1 mg/mL RNase A (Thermo Fisher Scientific, EN0531), and 10 U/µL RNase T1 (Thermo Fisher Scientific, EN0541) in 1× RNaseH buffer. The samples were then washed twice with PBST for 5 mins each. After the final PBST wash, the samples were incubated with 300 µL of splint ligation mixture containing 0.2 mg/mL BSA (New England Biolabs, B9000S), 2.5 µM splint ligation primer, and 0.1 U/µL T4 DNA ligase (Thermo Fisher Scientific, EL0011) in 1× T4 DNA ligase buffer at room temperature for 4 hours with gentle shaking. Subsequently, they were washed three times with 500 µL PBST for 5 mins each.

To create nanoballs of cDNA (amplicons) containing multiple copies of the original cDNA sequence, each cDNA circle undergoes linear amplification through rolling-circle amplification (RCA). This is achieved by immersing the cDNA in a 300 µL RCA mixture consisting of 0.2 U/µL Phi29 DNA polymerase (Thermo Fisher Scientific, EP0094), 250 µM dNTP, 40 µM 5-(3-aminoallyl)-dUTP, and 0.2

mg/mL BSA in 1× Phi29 buffer at 30°C for 4 hours with gentle shaking. Following RCA, the samples were subjected to two washes with PBST. Subsequently, they were incubated with 20 mM methacrylic acid N-hydroxysuccinimide ester (Sigma-Aldrich, 730300-1G) in 100 mM sodium bicarbonate buffer at room temperature for one hour, followed by two additional washes with PBST for 5 mins each. The samples then experience a 10-minute incubation in 500 µL monomer buffer containing 4% acrylamide (Bio-Rad, 161-0140) and 0.2% bis-acrylamide (Bio-Rad, 161-0142) in 2× SSC (Sigma- Aldrich, S6639) at 4°C. Following the aspiration of the buffer, a 35 µL polymerization mixture, made of 0.2% ammonium persulfate (Sigma-Aldrich, A3678) and 0.2% tetramethylethylenediamine (Sigma-Aldrich, T9281) dissolved in monomer buffer, is placed at the core of the sample and is promptly covered with a Gel Slick-coated coverslip. The polymerization is then carried out inside an $N_2$ enclosure for 90 mins at room temperature. Afterward, the sample is washed three times with PBST, each time for 5 mins.

Several iterative sequencing experiments were conducted to decode the rRNA identity. For each iteration, the sample initially underwent treatment with a stripping buffer containing 60% formamide (Calbiochem, 655206) and 0.1% Triton-X-100 (Sigma-Aldrich, 93443) at room temperature twice for 10 mins each, followed by a triple wash in PBST, each lasting 5 mins. Then the samples were incubated with a 300 µL sequencing mixture containing 0.2 U/µL T4 DNA ligase, 0.2 mg/ml BSA, 10 µM reading probe, and 5 µM fluorescent decoding oligos in 1× T4 DNA ligase buffer for at least 3 hours at room temperature. Post-incubation, the samples were thrice washed with a washing and imaging buffer made of 10% formamide in 2× SSC buffer, each wash lasting for 10 mins. Following the washing steps, the samples were immersed in the washing and imaging buffer for imaging. DAPI (Sigma-Aldrich, D9542) was dissolved in the wash and imaging buffer and performed following manufacturer's instruction for nuclei staining for 20 mins. Images were captured using a Leica TCS SP8 confocal microscope equipped with a 40× oil immersion objective (NA 1.3) and an acquisition voxel size of 142 nm × 142 nm × 500 nm.
